# Molecular dynamics study of stiffness and rupture of axonal membranes

**DOI:** 10.1101/2024.04.08.588595

**Authors:** Maryam Majdolhosseini, Svein Kleiven, Alessandra Villa

## Abstract

Diffuse axonal injury (DAI), characterized by widespread damage to axons throughout the brain, represents one of the most devastating and difficult-to-treat forms of traumatic brain injury. Different theories exist about the mechanism of DAI, among which one hypothesis states that membrane poration of the axons initiates DAI. To investigate the hypothesis, molecular models of axonal membranes, incorporating 25 different lipids distributed asymmetrically in the leaflets, were developed using a coarse-grain description and simulated using molecular dynamics techniques. Different protein concentrations were embedded inside the lipid bilayer to describe the different sub-cellular parts in myelinated and unmyelinated axons. The models were investigated in equilibration and under deformation to characterize the structural and mechanical properties of the membranes, and comparisons were made with other subcellular parts, particularly myelin. Employing a bottom-top approach, the results were coupled with a finite element model representing the axon at the cell level. The results indicate that pore formation in the node-of-Ranvier occurs at a lower rupture strain compared to other axolemma parts, whereas myelin poration exhibits the highest rupture strains among the investigated models. The observed rupture strain for the node-of-Ranvier aligns with experimental studies, indicating a threshold for injury at axonal strains exceeding 10 − 13%epending on the strain rate. The results indicate that the hypothesis suggesting mechanoporation triggers axonal injury cannot be dismissed, as this phenomenon occurs within the threshold of axonal injury.

**Highlights:**

- Developing a realistic molecular model of axolemma based on experimental data about its lipid composition
- Investigating how lipid composition and protein concentration affect the membrane’s structural and mechanical properties
- Identifying the most vulnerable regions of the axonal membrane

## 1. Introduction

Traumatic brain injury (TBI) is a brain pathology caused by an external force applied to the brain in traffic or sports accidents, falls, or violence [1]. Diffuse axonal injury (DAI) is a common type of TBI that affects the brain at cellular and sub-cellular levels. Swollen and disconnected axons are indicative of DAI at the cellular level [2, 3]. However, mild to low-moderate DAI cannot often be diagnosed by conventional imaging techniques, while these injuries cause microscopic pathologies at molecular level to the brain tissue that can disrupt normal brain function [4]. This makes it essential to investigate cellular and sub-cellular responses to external forces on the brain to gain more insights into the mechanism of mild DAI.

Axons are elongated projections of nerve cells. They can be myelinated or unmyelinated. Myelinated axons are wrapped by a multi-layer sheath called myelin. The myelin sheath covers the axon in segments, and gaps between the segments are called nodes of Ranvier. The axonal membrane or axolemma contains many membrane proteins that act as ion channels and transporters and play a vital role in the physiological functioning of cells. The concentration of these proteins is not uniform along the axon. This has been shown to affect the axonal membrane mechanical properties and their behaviour during an impact [5, 6]. The node-of-Ranvier exhibits a higher concentration of ion-channel proteins in the membrane, while unmyelinated axons have, on average, a lower ion-channel concentration [7]. Fig. 1 provides a schematic representation of a neural cell along with a sub-cellular representation of axonal membranes. Among the proteins present in the axolemma, voltage-gated sodium channels are one of the most abundant in the brain cells. In particular, the sodium channel, Nav1.1, is dominant in axon initial segments and the node-of-Ranvier [8, 9].

**Fig. 1:**
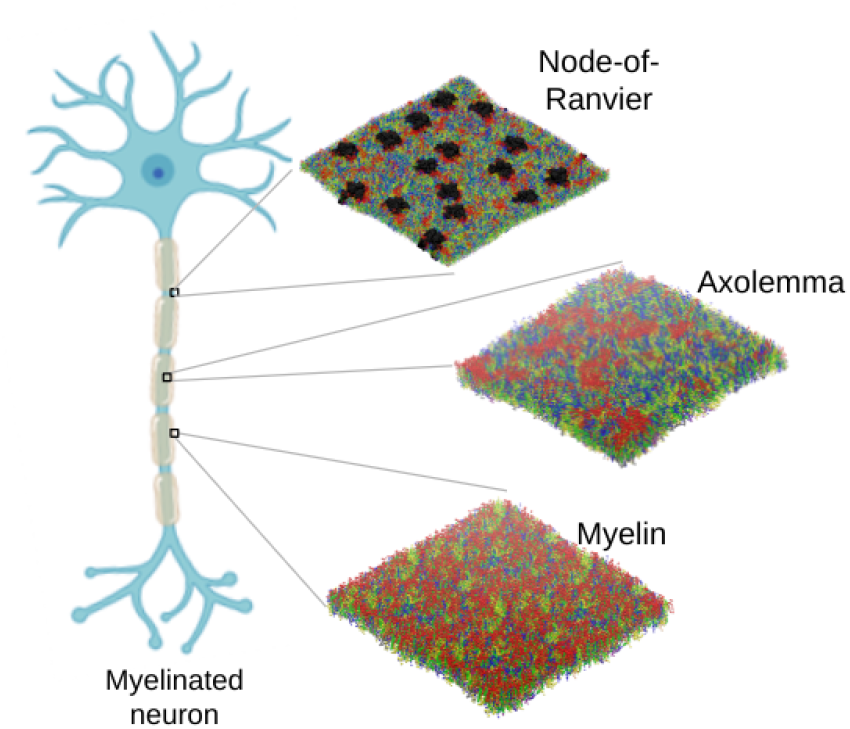
A schematic view of a myelinated neuron together with snapshot of the simulated bilayers. The arrow connects the bilayers to the corresponding element in the neural cell. Axolemma and the node-of-Ranvier have the same lipid composition. Lipids are illustrated as follows: PC lipids are in blue, PE in yellow, SM in orange, PS in purple, glycolipids in red and CHOL in green. The proteins are represented in black. Water and ions are not shown. Lipid head groups are abbreviated as follows: phosphatidylcholine (PC), phosphatidylethanolamine (PE), sphingomyelin (SM), phosphatidylserine (PS), and cholesterol (CHOL). The neuron model was taken from BioRender.com.

The lipid composition of membranes in different cells and subcellular elements varies significantly according to experiments. This can affect the function and properties of the cells (Table 1) [10]. Various studies have investigated how lipid content and cholesterol concentration affect the mechanical and structural properties of lipid bilayers [11, 12, 13, 14, 15, 16, 17]. These studies have highlighted not only the significant impact of molecular composition on membrane mechanical properties such as area compressibility [11, 12, 13, 14, 16], but also the effect on processes like mechanoporation [14, 15, 16, 17], interdigitation [13, 15], and bilayer thinning [12, 17]. The dependence of bilayer properties on lipid composition emphasizes the necessity for an accurate and specific molecular model to describe the axonal membrane. Capturing the unique composition of the axolemma will enable a more comprehensive investigation of the axolemma’s properties and behaviour under deformation.

**Table 1:** Experimental and modelled lipid composition (weight fraction) of different membranes. For experimental data, the mean weight fraction is provided along with its corresponding standard deviation. For abbreviations, see Fig. 1

|  | PC | PE | SM | PS | Glycolipid | CHOL | Other |
| --- | --- | --- | --- | --- | --- | --- | --- |
| **Experimental lipid composition (% weight)** |  |  |  |  |  |  |  |
| Axolemma<br>[18, 19, 20] | 21 ± 5 | 20 ± 5 | 3 ± 4 | 4 ± 2 | 20 ± 12 | 24 ± 3 | 4 ± 2 |
| Myelin [21] | 11 ± 1 | 17 ± 1 | 6 ± 3 | 6 ± 1 | 28 ± 3 | 28 ± 1 | 4 ± 2 |
| Brain white<br>matter [22] | 12 ± 1 | 20 ± 2 | - | 9 ± 1 | 24 ± 5 | 27 ± 6 | 9 ± 2 |
| **Molecular model lipid composition (% weight)** |  |  |  |  |  |  |  |
| Axolemma<br>model; De-<br>tails in section<br>2.1 | 22 | 23 | 3 | 4 | 15 | 29 | 5 |
| Myelin<br>model; De-<br>tails in [16] | 12 | 22 | 3 | 9 | 23 | 29 | 2 |
| Plasma mem-<br>brane model;<br>Details in [23] | 30 | 17 | 17 | 6 | 5 | 17 | 8 |

Investigations of axonal behaviour under deformation at cellular and sub-cellular levels have employed techniques such as atomic force microscopy [24], laser-induced cavitation [25, 26], transient pressurized fluid deflection [27], cell culture deformation [27, 28, 29, 30], Fluorescent Tracing [31], light microscopic examination [32] as well as computational modellings [15, 33, 34, 35]. The results of these studies have led to multiple hypotheses about the initial trigger of injury. One hypothesis regarding the primary mechanism of axonal injury is mechanoporation, which involves the formation of pores in the axonal membrane [29, 30, 35, 15, 31, 32, 25]. Another proposed hypothesis is that cytoskeletal damage is the initial trigger of axonal injury [24, 36, 27]. However, computational studies utilizing FE models of the axon, incorporating major cytoskeletal components and the axonal membrane, have questioned the hypothesis of cytoskeletal damage initiating axonal injury [35]. These FE studies have demonstrated high levels of strain primarily on the axonal membrane rather than the microtubules, suggesting that the membrane is more susceptible to injury than this cytoskeletal element. Capturing sub-cellular mechanisms of injuries and data on local strains and stresses from experiments is challenging due to their time and length scales that are in order of microseconds and nanometres. In addition, performing experiments on subcellular elements of axons is challenging due to their complex nature [37, 38]. Alternatively, to obtain more detailed data, modelling techniques like finite element (FE) modelling and molecular dynamics (MD) simulations can be combined to study brain injuries and clarify their mechanisms at sub-cellular and molecular levels.

Here, we aim to gain a better understanding of whether the mechanoporation of axonal membranes can trigger axonal injury using molecular modelling. To achieve this, we first need to develop a model that accurately describes the composition of the axolemma. Then, using MD techniques, we can tackle which sub-cellular compartment of the axon undergoes poration first and how lipid composition and protein concentration affect the mechanical properties of membranes. For this purpose, we have developed a coarse-grained (CG) model for axolemma based on experimental data. To elucidate the effect of the protein in the membrane, we embed different concentrations of Nav1.1 protein inside the bilayer. In total, we derived three models representing different parts of the axon. The models were simulated in equilibrium and under deformation. Finally, the systems were subjected to uniaxial deformation to simulate an injury condition. According to the hypothesis, membrane poration was considered as the time when axonal damage was initiated. Therefore, in the last step, we combined the deformation results in molecular simulations with the equivalent deformation of axolemma in an FE model of an axon at cellular level under uniaxial tension [39]. An overview of the workflow for simulations and analysis is provided in Fig. 2.

**Fig. 2:**
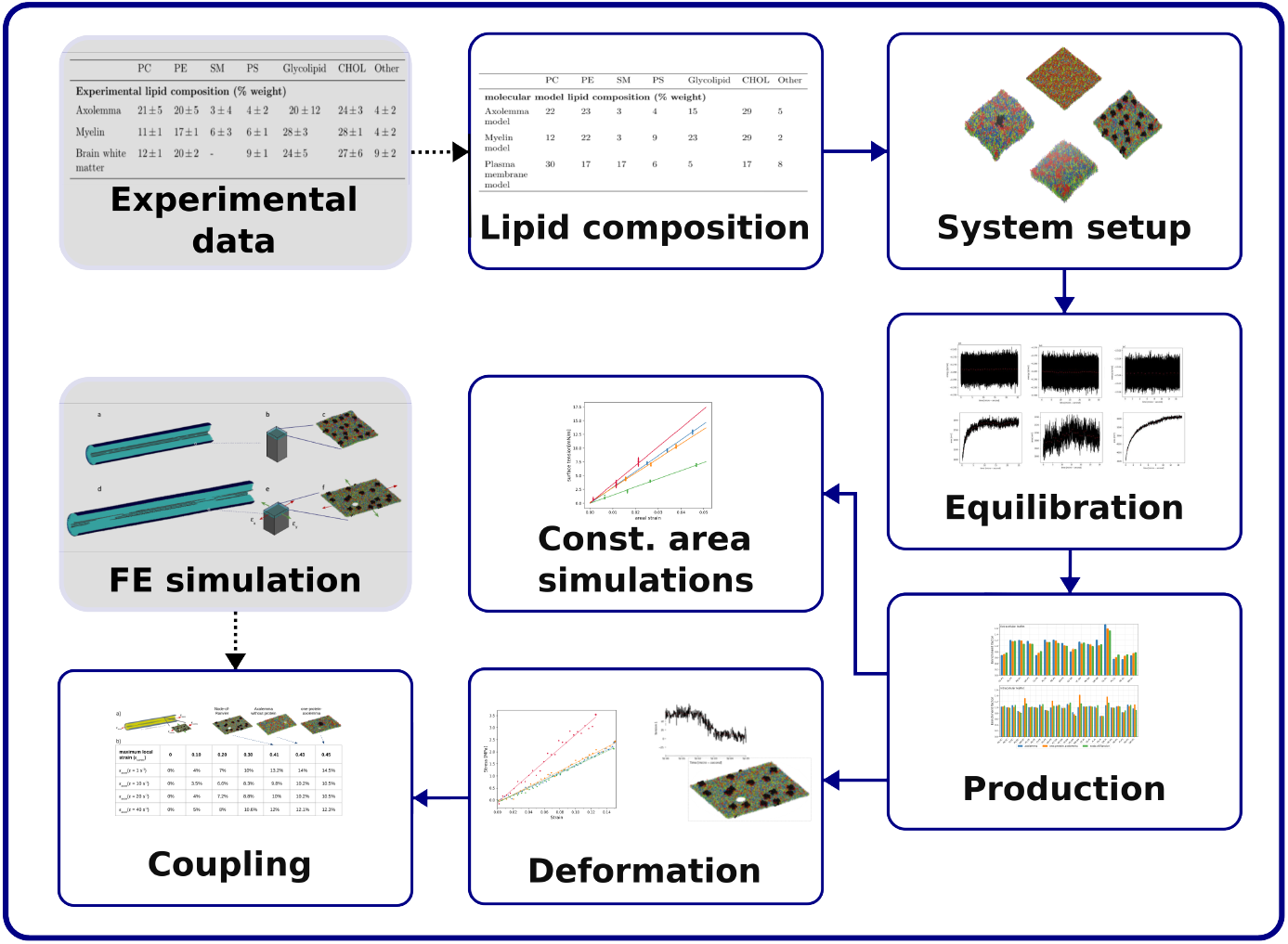
An overview figure about the steps taken for simulation and analysis of the systems

## 2. Methods

### 2.1. Molecular model for the axon

Axolemma is described as a bilayer consisting of different types of lipids distributed asymmetrically in the extracellular and cytoplasmic leaflets. We build the model utilizing available experimental data on the lipid composition for the axolemma, the lipid distribution within the extracellular and cytoplasmic leaflets, and the information on saturation index [19, 40, 41, 16]. Lipid-type composition and distribution between leaflets, saturation index, and tail length were determined using a combination of experimental data and information from established brain-type membrane models. While overall lipid composition of axolemma, fatty acid composition and unsaturation index of phosphatidylcholine (PC) and phosphatidylethanolamine (PE) were derived from experiments on human brain reported by DeVries et al. [19], neuron data reported by Fitzner et al. was used to derive chain length and saturation number in PS [40]. In case the experimental data on axolemma were missing, information on more general brain-type membrane models were used [41, 16]. Lipid disposition in the extracellular and cytoplasmic leaflet was derived from the brain membrane model developed by Ingólfsson et al. [41].

The final model is described as a bilayer consisting of 25 lipid types and cholesterol(CHOL), distributed asymmetrically in the extracellular and cytoplasmic leaflet of the axon cells. The bilayer composition (% mol) is 17% phosphatidylcholine (PC), 18% phosphatidylethanolamine (PE), 3% phosphatidylserine (PS), 4% phosphatidylinositol(PI), 2% sphingomyelin (SM), 17% glycolipids and 45% CHOL. The weight and mol fractions are reported in Table 1 together with experimental values. The specific lipid types used in developing the model, along with their mole percentage in each leaflet, are provided in Supplementary Table S1. The extracellular leaflet primarily consisted of 20% PC, 3% SM, 21% glycolipids, 44% CHOL and 1% other lipids, while the cytoplasmic leaflet contained 14% PC, 26% PE, 5% PS, 7% PI, 46% CHOL and 2% other lipids in % mol. The model had a lipid composition in line with the reported experimental values, except for a slightly higher weight fraction of CHOL in the model compared to the experimental value.

The tail length of the developed model varies between 16 to 24. Approximately 80% of the lipid tails in the axolemma are between 16 and 18 carbons in length in both extracellular and cytoplasmic leaflets. Approximately 90% of the lipid tails of PC, SM and PI have lengths of 16-18 carbons, while longer tail lengths of 20–24 carbons are present in about 54% of PS, 37% of PE and 30% of GL lipids. The saturation index of the axolemma model (ratio of saturated to unsaturated tails) is 0.47 in the extracellular leaflet and 0.42 in the cytoplasmic leaflet of the developed axolemma model.

The cryo-EM structure of human Nav1.1, as presented by Pan et al. [42] (PDB ID: 7DTD), was used as the starting structure for the protein system. Experimentally, the density of sodium channels in the axonal membrane differs across regions, ranging from 5 to 3000 *channels/µm*^2^, with the highest concentration found at the node-of-Ranvier [43, 7, 44]. Three models were built with different protein concentrations in the membrane to mimic different regions of the axon: a model without protein, called the “Axolemma” model, one with only one embedded protein (“one-protein-axolemma”) and one with 16 embedded proteins, with an equivalent protein concentration of 2519 *channels/µm*^2^, labelled as “node-of-Ranvier”. In addition, to compare the behaviour of the axolemma membranes with myelin, we use a previously developed and equilibrated CG model [16]. The model is implemented in Martini2.2 and can be downloaded from github(*https://github.com/alevil − gmx/myelin_m_odel)*. Table 1 shows the experimental and modelled lipid composition of the developed myelin model.

### 2.2. Molecular dynamics simulation

#### 2.2.1. System setup

All the molecules were described at the coarse-grained (CG) level using Martini 2.2 force field [45]. The starting structure of the membrane without the protein was built using INSANE (INSert membrANE) bilayer builder [46] and Martinizer python code was employed to transform the protein’s all-atom structure into a coarse-grained model [47]. The protein’s overall shape was constrained using the ElNeDyn approach [48], to control the overall conformation of the protein. A bilayer was first built using 1004 lipids and 820 cholesterol solvated by 52177 water beads, with 0.15 mol concentration of Na-Cl as ions in a box of size 20 × 20 × 20 *nm*^3^. Then, equilibration run was performed for the lipids to find their most suited neighboring configuration. The equilibrated system was then used to build a larger system 38 × 38 × 24 *nm*^3^. This system was equilibrated under the same condition for 20 microseconds before deformation.

In order to simulate the node-of-Ranvier, a larger concentration of proteins was embedded inside the membrane according to the following procedure. First, one protein was embedded in an equilibrated axonal membrane model of size 20 × 20 × 20 *nm*^3^ and the system was equilibrated again. Then, a larger system with a box size of 80 × 80 × 20 *nm*^3^ containing 16 proteins was developed from a previously equilibrated model, representing a system with a protein concentration of 2519 *channels/µm*^2^.

#### 2.2.2. Equilibration and deformation

All the simulations were performed in the GROMACS simulation package[49], version 2020 and 2021 [50, 51]. After energy minimization, all the systems were equilibrated at NPT condition. The pressure and temperature were kept constant at 1 bar using C-rescale barostat [52] (*τ*_*p*_ = 12.0 *ps, κ* = 3 × 10^−4^ *bar*^−1^) and at 310 K using V-rescale thermostat [53] (*τ*_*T*_ = 1.0 *ps)*, respectively. The time-step was 10 *f s* with Verlet cut-off scheme for non-bonded interactions that were updated every 20 steps with a cutoff distance of 1.1 nm for both Coulomb and van der Waals interactions. Long-range interactions were defined using reaction field. The models containing proteins were first equilibrated by having position restraints on the proteins to relax lipid distribution around the protein and then, the position restraints were removed to allow the proteins to move freely as well. For the one-protein-axolemma, the protein-lipid contacts and energy of the system did not change much after removing the position restraint of the protein (supplementary Fig. S 1); therefore, the last frame of the system with restrain on the protein was used to run the deformation simulations. However, the proteins in the node-of-Ranvier model tend to move when they are free, so the deformation is applied to the system after being equilibrated for 11 *µs* with no position restraint. Table 2 provides more information about each system, their structure, and performed MD simulations, while more detailed data about developing the systems are provided in the supplementary Table S2.

**Table 2:** Information on performed molecular dynamics simulations.

| Model Name | Dimension<br>(nm) | # Nav1.1<br>proteins | Equilibration<br>time ($\mu s$) | No. of replicas<br>at each<br>deformation<br>rate ($\frac{nm}{ps}$) | | |
| --- | --- | --- | --- | --- | --- | --- |
| | | | | $10^{-5}$ | $10^{-6}$ | $10^{-7}$ |
| Axolemma (small) | $20 \times 20 \times 20$ | 0 | 25 | No deformation | | |
| Axolemma | $38 \times 38 \times 24$ | 0 | 20 | 1 | 3 | 3 |
| One-protein-axolemma | $46 \times 38 \times 19$ | 1 | $30^a$ | 1 | 3 | 3 |
| Node-of-Ranvier | $80 \times 80 \times 20$ | 16 | $15^a + 11.5$ | 1 | 3 | 3 |
| Myelin | $41 \times 41 \times 10$ | 0 | by [16] | 1 | 3 | 3 |

<sup>a</sup>: simulation performed by applying position restraint on the protein

To investigate pore formation under strain, we initially deformed the systems at a rate of 1×10^−5^ *nm/ps* in the x-direction until a 30% deformation was achieved. Following this, two additional deformations were applied to the pre-deformed models at slower rates of 1 ×10^−6^ *nm/ps* and 1 × 10^−7^ *nm/ps*. This approach enabled us to analyze the effects of deformation rate on pore formation. Each simulation was conducted at least three times. For comparison, similar deformation protocols were applied to the equilibrated myelin model described by Saeedimasine et al. [16], allowing us to evaluate the properties of the developed systems relative to the myelin structure.

All the simulations were conducted on the Dardel supercomputer, where each compute node is equipped with two AMD EPYC™ Zen2 2.25 GHz 64-core processors, offering a total of 128 physical CPU cores per node. A single node achieves a performance of 1000 ns/day for the axolemma system (approximately 300,000 particles). This means that, on average, we require 10,400 node hours (54 days) to obtain results for each of the four molecular systems, from equilibration to pore formation. Consequently, this amounts to more than 5.4 million CPU hours of simulation time. Additionally, it is worth noting that GROMACS is highly efficient at running parallel simulations and multiple CPUs can be used in parallel to run the simulations [49].

To check the convergence of the simulations at equilibration, total energy and surface area of the bilayers were calculated over time. Both properties reached equilibrium within the first 10 *µs* for all models (supplementary Fig. S 2). In addition, lipid distribution in the bilayer in the axolemma model was checked and compared with the small axolemma system in the last 10 microseconds by calculating the lipid neighbouring. We assumed that the system reached equilibrium when the average number of lipid neighbouring along with the surface area of the bilayer and energy of the system converged (see supplementary Fig. S 3).

### 2.3. Structural and mechanical properties

#### 2.3.1. System in equilibration

In order to check the equilibration of the system and the lipid aggregation in equilibrium, enrichment factor was calculated and compared for the last 10 *µs* of equilibrium for each lipid type based on the method proposed by Corradi et al. [54]:

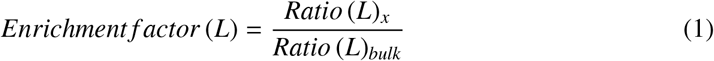

In which: *Ratio*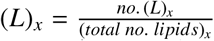, is calculated by dividing the number of type L lipids within the distance of x from the given lipid by total number of lipids within the distance of x from the given lipid and *Ratio*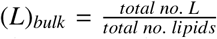, is defined as the total number of type L lipids in the system divided by total number of lipids.

For the systems with protein, we use *Ratio* (*L*)_*x*_ to define the protein neighboring. The cut-off distance was set to be 1.5 and 2.1 nm to calculate lthe ipid-lipid enrichment factor in athe xolemma model and protein-lipid nneighbouringin one-protein-axolemma and the node-of-Ranvier, respectively, in line with the distance chosen by Saeedimasine et al. [16].

#### 2.3.2. System under deformation

To derive the mechanical properties of the systems, such as Young’s modulus, principal stress in x direction, *σ*_*x*_, and strain, *ϵ*_*x*_, values were calculated as follows:

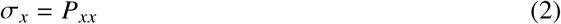

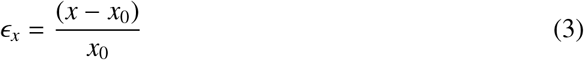

Where*P*_*xx*_ is the lateral pressure in the x direction, the same direction from which the system is deformed [55], *x* is the bilayer dimension in x direction at different deformation points and *x*_0_ is defined as the x dimension of the bilayer when the average surface tension of the system is zero. Values of *P*_*xx*_ and *x* are averaged at time-windows of 0.1 µ*s*.

The stress-strain curve of a material under deformation acts differently at different stages of deformation. At small strains, called the elastic region, there is a linear relationship between stress and strain curve, and the slope of the line in this part is representative of the stiffness or Young’s modulus of the system. The strain in which the stress-strain curve becomes non-linear depends on the system and its mechanical properties. The elastic region for the simulated systems in this study was until the deformation of 15%. Equation 4 shows how Young’s modulus is calculated in the elastic region.

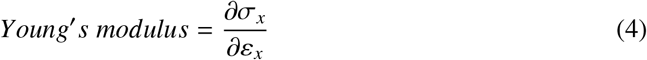

in which *σ*_*x*_ is the uniaxial stress, and *ε*_*x*_ is the strain of the system. The Young’s modulus for the systems is derived from the stress-strain curve of the systems under deformation with the strain rate of 1 ×10^−5^ *nm/ps* by employing the ordinary least squares regression method in Python to fit a line to the strain-stress curves and the slope of the line along with the standard error of the fit are presented.

Another mechanical property of the bilayer is the area compressibility modulus, *K*_*A*_, which is defined as the derivative of the surface tension *γ* with respect to areal strain *ε*_*A*_ at constant temperature, experimentally [56, 57].

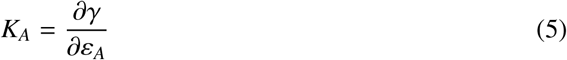

Assuming that the surface area of the bilayer is equal to the surface area of the box, the areal strain was derived by dividing the area of the box at different deformations to the initial area of the box at strain of zero 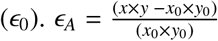. Surface tension can also be calculated as:

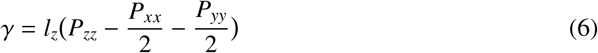

in which *P*_*xx*_, *P*_*yy*_ are the lateral pressure, *P*_*zz*_ is the normal pressure and *l*_*z*_ is the height of the box.

At small deformations, there is a linear relationship between surface tension and areal strain and consequently, area compressibility modulus is constant. Therefore, to calculate this parameter, we simulated the systems in constant pressure and areal strain condition, *NP*_*z*_*AT*, in small areal strains of up to 4.5% for 6 *µs* and using the data of the last 4 *µs* of simulations to make sure the system is in equilibrium, the production data was processed every 1 *µs* and the average surface tension, *γ*, and strain values were used to calculate the area compressibility modulus to make the results more comparable with experimental data. *K*_*A*_ is calculated using the same regression method used to calculate Young’s modulus. Using strain-stress curves calculated from these simulations, we also calculated Young’s modulus of each system for comparison of the results with stiffness value derived from the stress-strain curves of the models under deformation.

Bilayer rupture is characterized by the formation of a pore in the bilayer under deformation, accompanied by a sudden decrease in surface tension. The strain at this point is defined as the rupture strain, *ϵ*_*pore*_. For each system, the average surface tension and its standard deviation were calculated 0.1 *µs* prior to pore formation. Additionally, the failure stress—corresponding to the average stress immediately before pore formation—was computed over the same time interval. This value represents the ultimate stress the membrane can withstand, providing insights into the maximum stress each system can endure before rupture.

### 2.4. Multi-scale approach

To investigate the membrane poration and how it leads to axonal damage at cellular level, we have coupled the molecular results with a generic FE model of the axon under stretch. The FE axon model was previously developed by Montanino and Kleiven and described in detail in Ref [35]. Briefly, the axon is modelled as an 8 *µm* cylinder consisting of microtubules, neurofilaments and tau proteins connecting the fibres together and an axolemma-cortex compartment surrounding the whole model. Periodic boundary conditions were applied to the neurite’s axonal direction. To account for the variability of microtubule discontinuity on axonal behaviour, 5 different axonal models with different microtubule discontinuity locations were developed and stretched. The axon is then subjected to deformation under uniaxial strain at different rates.

The strain applied to the axon leads to non-uniform deformation across its sub-cellular components. Employing a bottom-up approach, we assume that each element of the cortex corresponds to the membrane systems developed at the molecular level. Therefore, the strain of the MD scale membrane at each point when deformed corresponds to the local strain of one element of the cortex in the FE model. Fig. 3 shows how the results from the FE and MD models are coupled.

**Fig. 3:**
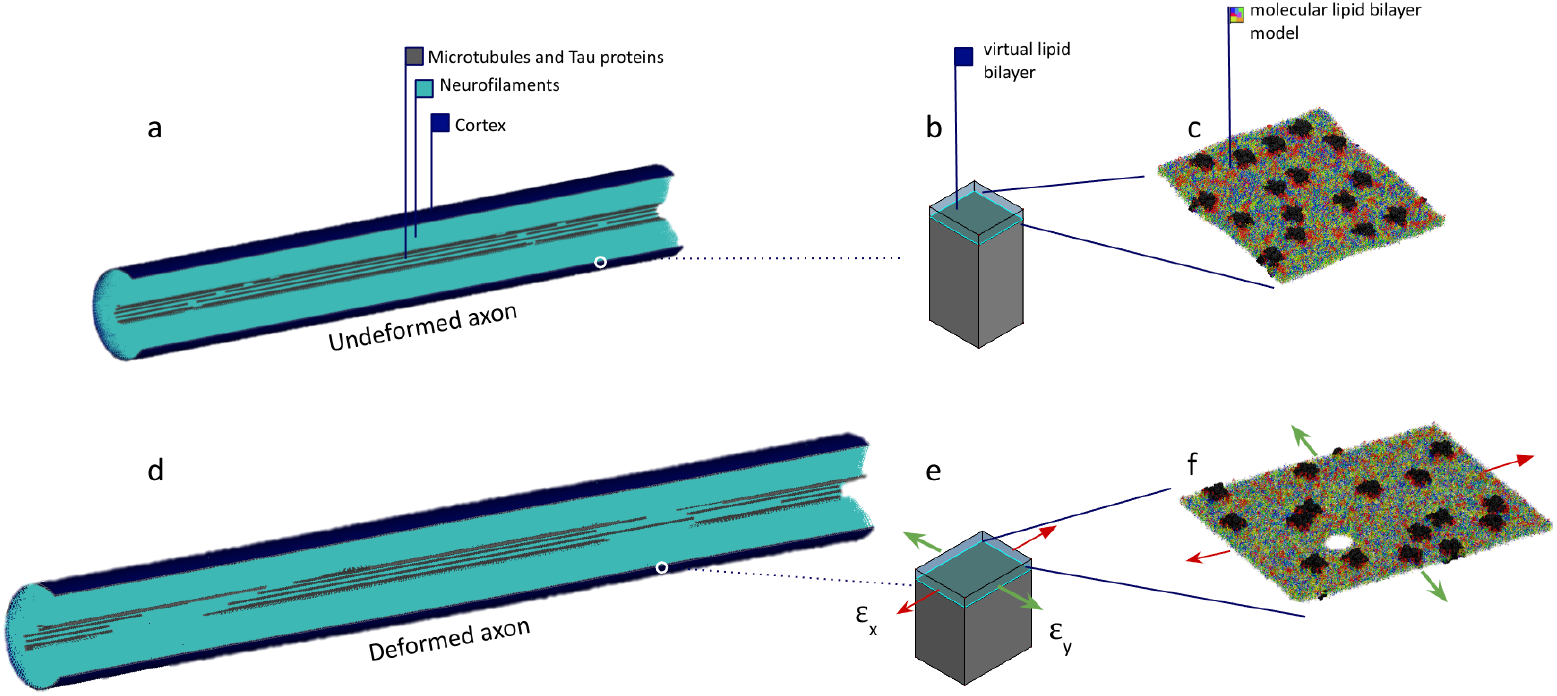
Coupling FE model of the axon in cellular level with molecular systems representing the axonal membrane [15]. a) FE model of the undeformed axon in cellular level; A quarter of the axon model is removed for better representation of the sub-cellular elements. b) A single element on the cortex in an undeformed state corresponding with the membrane models in molecular level. c) Undeformed lipid bilayer of the axolemma with one embedded protein. d) FE model of the axon under uniaxial deformation in cellular level. e) The cortex element with the maximum local strain during the axonal deformation is extracted. f) Lipid bilayer of the axolemma with one embedded protein undergoing the maximum local strains of the cortex element (figure adapted and modified from [15]).

To calculate the axonal strain at which membrane poration occurs, we identified the cortex element with the highest local strain and mapped those values to the obtained total axonal strain. Furthermore, the strain in which the mechanoporation occurs at molecular level when the model is deformed is assumed to be the threshold of the axonal injury at cellular level.

## 3. Results

The membrane models have been simulated at equilibrium as well as under deformation to mimic accidental TBI. The structural properties at equilibrium are presented in section 3.1 while the mechanical properties of the membranes under deformation, including the rupture strain, and the coupling of the results with FE models are discussed in section 3.2.

### 3.1. Structure of the axolemma membranes

The axolemma model was built by combining experimental data from different sources on head group fractions, tail lengths and saturation index since the complete lipidome for the axolemma is not experimentally available to the best of knowledge. The resulting bilayer composition agrees well with the average experimental lipid composition (Table 1), but we can expect some deviations because of a lack of experimental information on the one-to-one relation between head groups, tail lengths and saturation index. The axolemma model differs in lipid composition not only compared to an average plasma membrane: it has a high weight fraction of CHOL followed by PC and PE phospholipids (see Table 1). A high CHOL content in the axolemma is expected since membranes of the brain cells are usually more enriched in CHOL compared with other cell membranes in the body [58, 59]. The current axolemma model is more enriched in PC lipids, less in glycolipids and CHOL than the myelin and average brain white matter membranes[16, 41]. In comparison with the plasma membrane, the axolemma model has a higher content of glycolipids and a lower content of PC and SM.

Lipid tails also influence the structural and mechanical properties of membranes. While longer tail lengths increase the thickness of the membranes [60, 61], fully saturated lipid tails not only increase the thickness but also affect the membrane rigidity [60, 61]. In addition, lipid tail mismatch can increase membrane compressibility [62]. The tail length of the axolemma is similar to the plasma membrane models, with 80% of short tail lengths of 16-18 in both extracellular and cytoplasmic leaflets. While the myelin has 48% short tails in cytoplasmic leaflet and 95% in the extracellular leaflet.

Here, we describe the membrane models that were designed for this work. For a detailed structure description of the myelin model at equilibrium, we refer to [16]. Lipid distribution for axolemma models was investigated by calculating the enrichment factor for each lipid type. The results shown in Fig. 4 suggest that the distribution of lipids inside the membranes is not uniform. However, all models follow the same trend of lipid aggregation and depletion regardless of whether a protein is embedded inside. While phospholipids in the extracellular leaflet tend to aggregate with each other, they tend to deplete the glycolipids. The same trend can be seen for glycolipids that aggregate most with each other. In the cytoplasmic leaflet where no glycolipid exists, lipids are distributed more uniformly; however, the PS lipids tend to deplete each other.

**Fig. 4:**
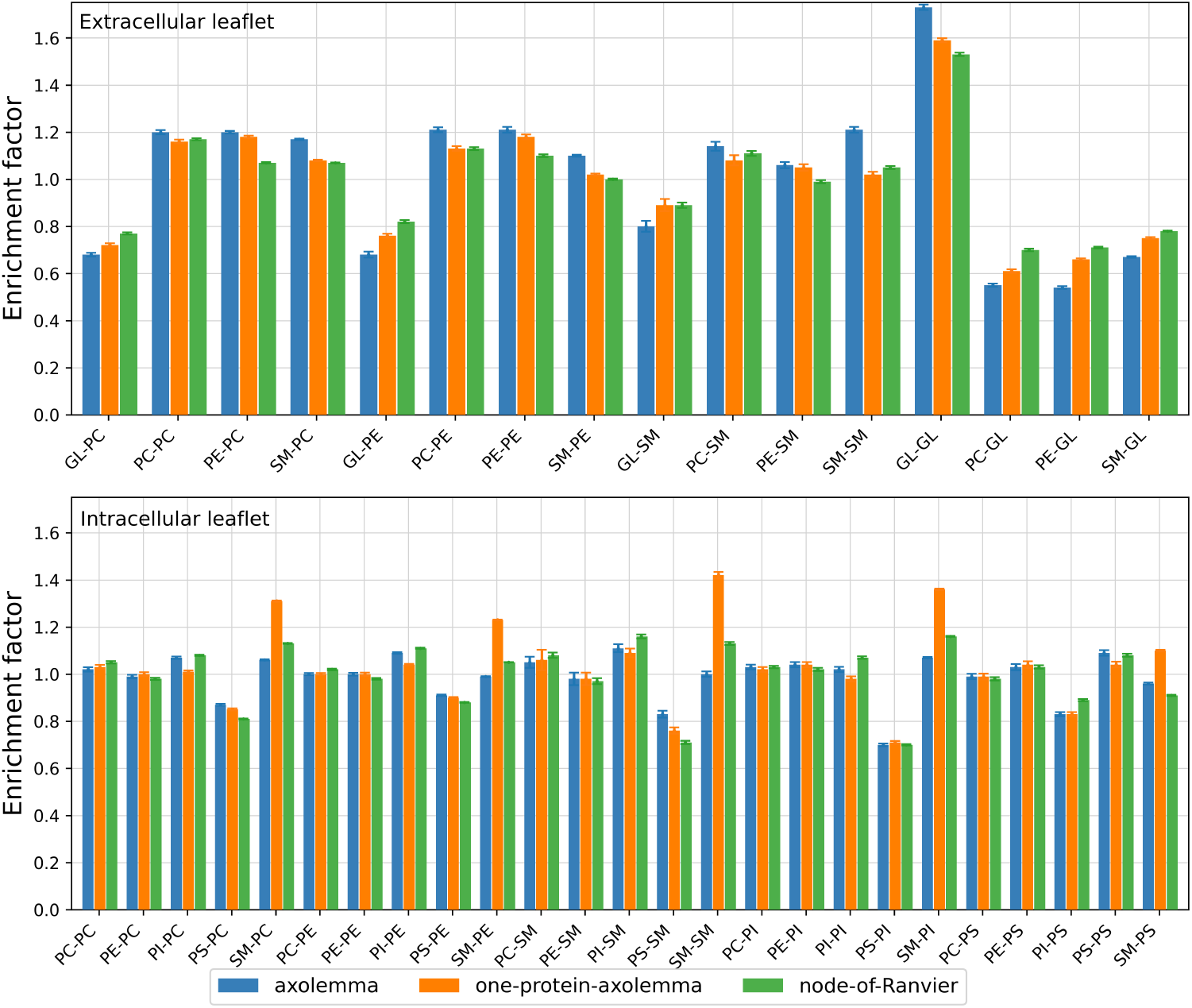
Average Lipid-Lipid enrichment of different systems in the last 10 *µs* of equilibration along with their standard deviation shown as error bars. For abbreviations, see Table 1

In the presence of proteins, the aggregation of glycolipids decreases in the extracellular leaflet. This change may be attributed to the fact that these lipids aggregate more with proteins, and as a consequence, the total lipids in their neighbouring decreases. In the cytoplasmic leaflet, SM lipids tend to aggregate more with other lipids in the one-protein-axolemma compared to the axolemma and the node-of-Ranvier models. However, as the number of SM lipids in the system is not much (only 2% of the composition), this change will not affect the properties of the system. Comparing the results with other models, it can be seen that the model of an average brain membrane developed by Ingolfsson et al. [41] also has a similar trend in the extracellular leaflet where the glycolipids co-localize and other lipids tend to aggregate with each other and deplete the glycolipids. Similar behaviour can be seen for the glycolipids in the model of the plasma membrane [23], but in this model, other lipids also aggregate with glycolipids, and less localization of the glycolipids can be seen. In the cytoplasmic leaflet, the distribution of lipids in the plasma membrane and brain membrane models [23, 41] are similar to the axolemma; however, the depletion of PS lipids is not seen in these models.

For the one-protein-axolemma model and the node-of-Ranvier, the lipid neighbouring around the proteins was also calculated, and the results are presented in Table 3. From the lipid-protein aggregation data in this table, it is indicated that in both systems, about half of the lipids surrounding the protein are glycolipids in the extracellular leaflet, while SM lipids tend to be depleted from the proteins. In the cytoplasmic leaflet where no glycolipid is present, PE phospholipids are more aggregated around the protein, while SM lipids deplete from the protein. Aggregation of glycolipids in the extracellular leaflet and PE phospholipids in the cytoplasmic leaflet along with SM depletion are also reported by Saeedimasine et al. [16] for the neighbouring of two different plasma membrane models with different protein concentrations.

**Table 3:** Protein-lipid neighboring for the axolemma model with protein and the node-of-Ranvier at the last 10 *µs* of equilibrium.

| System | Extracellular leaflet |  |  |  | Cytoplasmic leaflet |  |  |  |  |
| --- | --- | --- | --- | --- | --- | --- | --- | --- | --- |
|  | PC | PE | SM | GL | PC | PE | SM | PI | PS |
| one-protein-axolemma | 0.25 | 0.15 | 0.02 | 0.57 | 0.19 | 0.49 | 0.01 | 0.13 | 0.18 |
| node-of-Ranvier | 0.27 | 0.18 | 0.04 | 0.51 | 0.21 | 0.47 | 0.02 | 0.14 | 0.16 |

### 3.2. Mechanical properties

To investigate the behaviour of the models during injury, the bilayers were deformed at different strain rates in *x* direction (Table 2 and supplementary Table S2). Based on the results, the calculated stiffness of the bilayer, rupture strain and other mechanical properties of the bilayer under deformation are reported in this section.

*Young’s modulus*. The values of Young’s modulus representing the stiffness of the bilayers are reported in Table 4 and the stress-strain points used for the fitting are plotted in Fig. 5. The results show that all the models with axolemma composition have similar stiffness of 15-16 MPa, while the myelin model has higher stiffness (26 MPa). This suggests that the protein concentration has little effect on the stiffness of the system, and this value is largely influenced by the lipid composition. Moreover, the stiffness values calculated from simulations performed at constant areal strains are 16.1, 14.9, 7.1 and 25.1 MPa for axolemma, one-protein-axolemma, node-of-Ranvier and myelin, respectively. While all the values are close to those derived from the stress-strain curve of the system under deformation, Young’s modulus derived for node-of-Ranvier is twice lower when calculated using the latter approach. The difference is because all systems show a stiffening behaviour by increasing the strain that is similar to the behaviour of biological soft tissues in toe-region. Although all models reach the linear region at around 2% strain, the toe region for the node-of-Ranvier lasts longer and the linear region starts at around 4%.

**Table 4:** Mechanical properties of different systems under deformation, the average rupture strain and surface tension 0.1*µs* before the pore for different tests are provided along with their standard error.

| Model | Rupture strain | Average surface tension before pore ($mN/m$) | Average stress before pore (MPa) | Young's modulus (MPa) | $K_A$ (mN/m) |
| --- | --- | --- | --- | --- | --- |
| Axolemma | $0.51 \pm 0.02^b$;<br>$0.43 \pm 0.03^c$ | $73.1 \pm 0.5^b$ ;<br>$70.1 \pm 0.6^c$ | $5.63 \pm 0.04^b$;<br>$5.05 \pm 0.03^c$ | $15.5 \pm 0.99^a$ | $289 \pm 7$ |
| One-protein-axolemma | $0.50 \pm 0.01^b$;<br>$0.45 \pm 0.01^c$ | $73.5 \pm 0.4^b$ ;<br>$71.4 \pm 0.4^c$ | $5.56 \pm 0.03^b$;<br>$5.19 \pm 0.03^c$ | $16.4 \pm 0.99^a$ | $266 \pm 8$ |
| Node-of-Ranvier | $0.44 \pm 0.01^b$;<br>$0.41 \pm 0.01^c$ | $72.3 \pm 0.2^b$ ;<br>$70.2 \pm 0.2^c$ | $4.84 \pm 0.02^b$;<br>$4.60 \pm 0.02^c$ | $15.3 \pm 0.99^a$ | $148 \pm 10$ |
| Myelin<sup>d</sup> | $0.53 \pm 0.07^b$;<br>$0.53 \pm 0.04^c$ | $76.8 \pm 0.78^b$ ;<br>$77.5 \pm 0.32^c$ | $8.19 \pm 0.05^b$;<br>$8.24 \pm 0.04^c$ | $28.1 \pm 0.98^a$ | $341 \pm 17$ |

\* Data calculated at different deformation rates of: a: $10^{-5}$ ; b: $10^{-6}$ ; c: $10^{-7} nm/ps$
d: for details on the myelin membrane model, see [16]

**Fig. 5:**
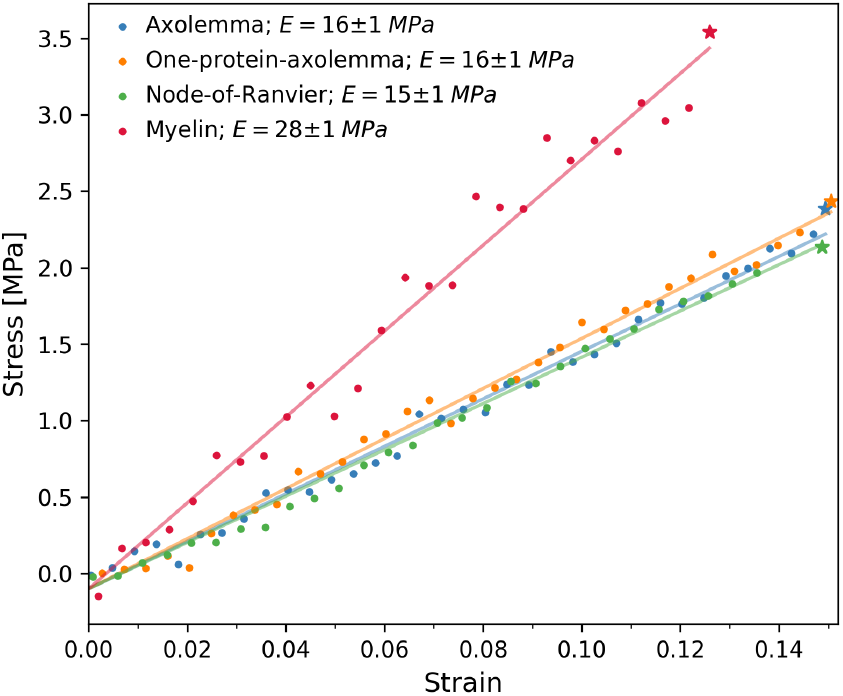
Strain Stress points of the systems in the elastic part of deformation and the corresponding fitted line using linear regression method. The stress value for each system at the end of the linear part representing the yield stress is shown by stars.

A direct comparison between our results and experiments is not possible, as to the best of our knowledge, no experimental data has measured the value of Young’s modulus for the axonal membranes under uniaxial deformation. Nevertheless, we try to compare the results with the stiffness of other lipid bilayers. Jaing et al. performed tensile tests on two different silicone bilayers [63] and stress-strain data until 200% were reported. We calculated Young’s modulus of the membranes based on the reported data for the first 15% of deformation using the linear regression method and derived the stiffness of 30 ±1 and 37± 2 MPa for these bilayers. Other studies applying compression to lipid bilayers using atomic force microscopy calculate a Young’s modulus of 19.3 to 28.1 MPa for a bilayer of DOPC/DPPC lipids [64], 27 to 31 MPa for POPC and PDPC bilayers [65] and 50 MPa for artificial lipid bilayers of DPhPC lipids [66]. The calculated Young’s modulus in the current study is in the corridor of the data derived for other membranes and lipid bilayers.

#### Compressibility modulus

Another mechanical parameter representing the response of the bilayer under tension is the areal compressibility modulus, *K*_*A*_. This value was calculated for the systems at three different areal strain values of 0.1 to 4.5%. The average surface tension along with standard errors in each *µs* is plotted as a function of areal strain in Fig. 6 and the results for the calculated areal compressibility modulus and the corresponding standard errors are reported in Table 4. Despite Young’s modulus that was quite similar for all the developed models of axolemma, *K*_*A*_ is affected by the concentration of proteins, as the node-of-Ranvier has almost half the compressibility modulus of the axolemma model with protein. Similar results were reported in previous studies on the model of plasma membrane [16]. The myelin has the highest compressibility modulus compared to other systems. The reported experimental values for *K*_*A*_ are from 179 to 450 mN/m for red blood cell membrane at 25 °C [56, 67, 68] which are in the range of values reported in this study. However, no experimental data is available for the area compressibility modulus of axonal membranes.

**Fig. 6:**
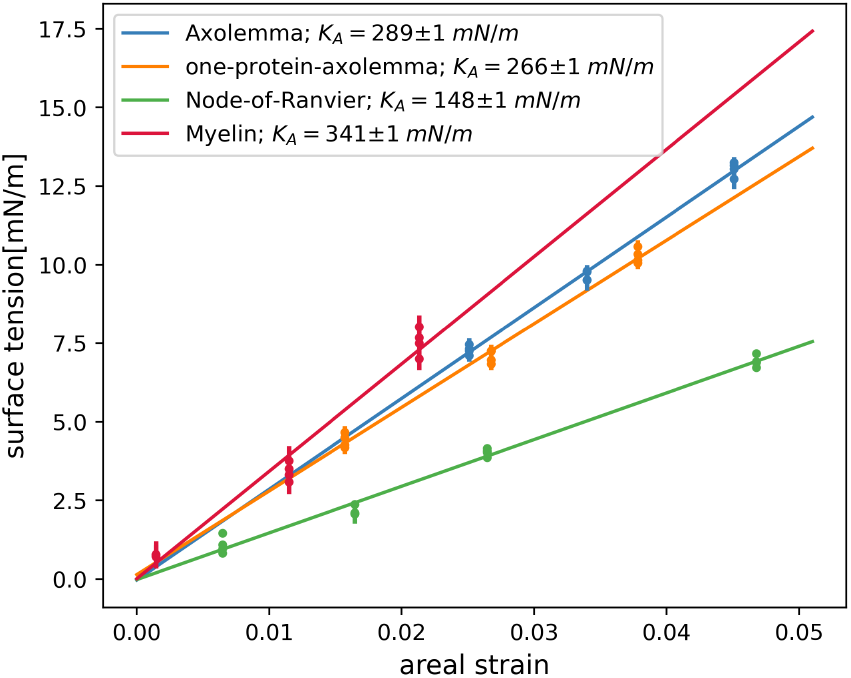
Values of surface tension (*mN/m*) and the corresponding areal strain of different systems. Data is reported as the average and standard error of the surface tension vs areal strain in time windows of 1 *µs* for each system when equilibrated in constant areal strain. Areal compressibility modulus of the systems is calculated as the slope of the line fitted to the data.

#### Membrane rupture

Assuming that the pore formation of the axolemma triggers the axonal injury, we deformed the systems until rupture at two different strain rates. The strain in which the rupture occurs depends on both the molecular system and the deformation rate. Table 4 displays the mean rupture strain and standard deviation across various replicas for each system at different deformation rates. The node-of-Ranvier model exhibits a lower rupture strain compared to other systems, while myelin has the highest rupture strain. The rupture strains in axolemma and one-protein-axolemma are close, with slightly higher rupture strain than the node-of-Ranvier.

The average surface tension before pore formation remains relatively similar for the axolemma models, both with and without proteins, across various strain rates. However, the average stress before pore formation increases proportionally with the rupture strain. The reported values for myelin surpass those for axolemma models due to myelin’s higher stiffness, higher *K*_*A*_, and higher rupture strain.

Finally, the threshold of injury at cellular level is defined by coupling the molecular results with the FE model of the axon under stretch. Fig. 7 illustrates the maximum local strain within the cortex in the FE model alongside the corresponding axonal strain under various strain rates. The strain where the mechanoporation occurs in MD is marked with the corresponding membrane. The results show that the node-of-Ranvier has a rupture strain of 41%, which corresponds to an axonal strain of 9.8 to 13.2 % depending on the strain rate, while the myelin seems to form rupture at higher strains compared to other membranes.

**Fig. 7:**
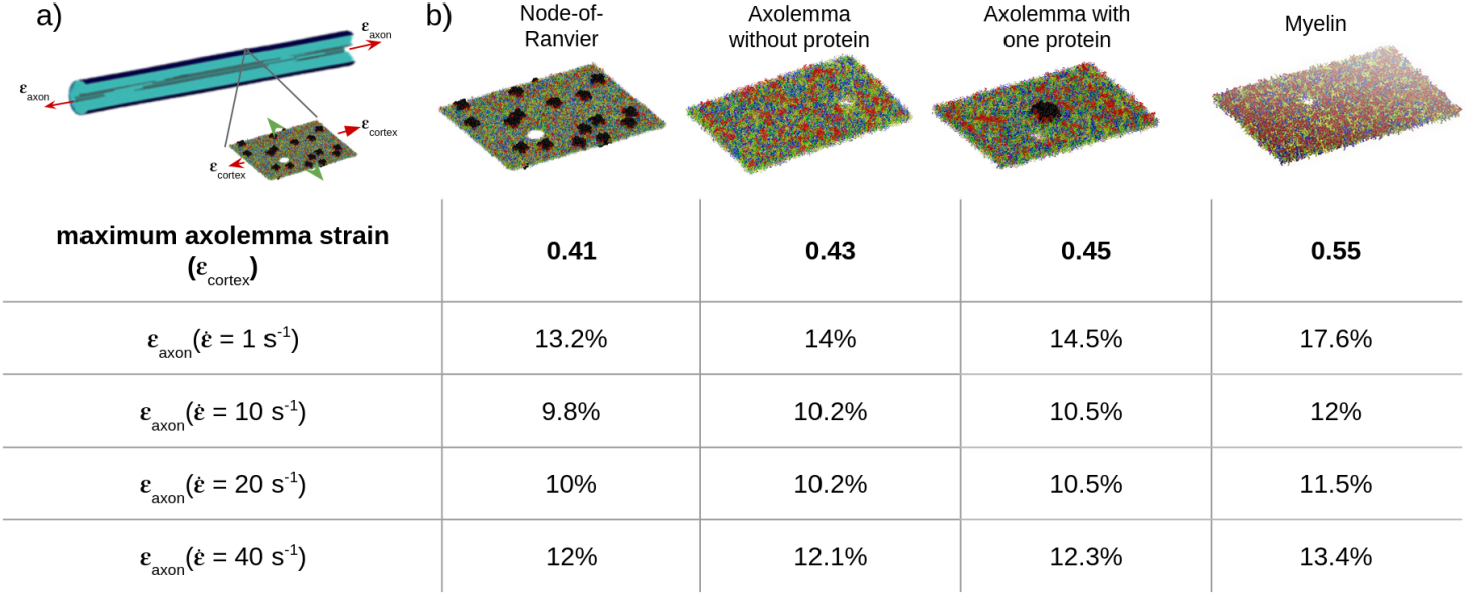
Coupling the molecular dynamics results with finite element model of the axon under deformation. a) A schematic view of deformation in the axon that leads to mechanoporation of the membrane and axonal injury. b) Axonal strain with the corresponding element with maximum local strain for different deformation rates are provided in the table, and the strain in which each system undergoes mechanoporation is marked.

As the developed axon model is based on an unmyelinated axon, we need to check the effect of the material properties of the cortex on maximum local deformation value and location. Therefore, we ran two separate FE simulations of the uniaxial strain of the axon with a rate of 10/s and modified cortex material properties: one to represent myelin and the other to represent the node-of-Ranvier. We assumed that the material properties change proportional to the compressibility modulus of the membranes at the molecular level. Therefore, we changed the material properties of the cortex in the FE model with the same proportion, assuming that cortex properties used in the literature represent the one-protein-axolemma model in our MD simulations. The results indicate that the maximum local strain occurs in similar elements across these models, with only minor changes in local strain values. Further details on the mechanical properties assigned to each model are provided in the supplementary table S3 and the maximum local strains for each model are presented in supplementary Fig. S 4

## 4. Discussion

Clinically, DAI can initially go undetected, with diagnosis often occurring days or weeks post-injury, as symptoms gradually emerge due to the progressive nature of this condition [69]. However, early diagnosis is vital for optimizing treatment, improving recovery outcomes, and enhancing patient care [70, 71]. Therefore, understanding the exact mechanisms that initiate DAI is essential for early detection of the injury. Previous research has indicated that initial signs of DAI post-impact include calcium (Ca^2+^) influx and focal axonal swelling, potentially resulting from membrane rupture. In the current study, we explored the hypothesis stemming from these findings, stating that mechanoporation of the axonal membrane acts as a trigger for axonal injury. To investigate this hypothesis, we used a multi-scale approach in which we used models of axons at cell level developed using FE analysis [15], while models of axonal membranes at molecular level were investigated using MD. Employing a bottom-top approach, the results from these two scales were coupled. The mechanoporation of the membrane models in MD was used to predict the threshold of axonal injury at cellular level.

Using experimental data about the lipid composition of axonal membranes and distribution of different lipid types in the cytoplasmic and extracellular leaflets, we developed a realistic axonal membrane model for a more accurate assessment of the rupture strain during injurious events. Previous studies have shown that membrane rupture strains and other mechanical properties, such as the area compressibility modulus and bending rigidity, are significantly influenced by lipid types and their distribution between the leaflets [72]. The asymmetric distribution of lipids between the leaflets also contributes to membrane curvature and influences interactions with proteins; namely, the distribution of charged lipids affects the electrostatic interactions with membrane proteins [73, 74, 75, 76].

The stiffness of lipid membranes is affected by structural differences like lipid composition, cholesterol content, and protein interactions. Previous studies have shown that membranes with high levels of saturated lipids tend to be more rigid, while unsaturated lipids lead to less tightly packed, more fluid, and thus softer membranes [77]. Membrane rigidity is also affected by cholesterol content: CHOL enhances lipid packing, reduces the fluidity of the bilayer, and restricts lipid movement [78, 79]. This is because of the rigidity of the sterol ring, which reduces lipid chain entropy and fluidity. This effect is more evident in unsaturated lipid chains [80]. In the current study, myelin shows a higher stiffness despite having more unsaturated lipids (72%)than the axolemma (69%). It might be due to the high cholesterol content in the myelin that affects the fluidity of unsaturated lipids, leading to a stiffer membrane. Furthermore, the presence of protein or protein concentration does not seem to affect the stiffness of the membrane. However, higher protein concentration reduced the compressibility modulus of the node-of-Ranvier, which can stem from the interactions of lipids with proteins. It has been shown that the protein modulates stiffness by inducing order or disorder in surrounding lipids, depending on the protein structure and the nature of lipid-protein interactions. For example, while some proteins enhance rigidity by increasing surface pressure and order within lipid domains [81], some reduce membrane stiffness by disrupting lipid packing and increasing membrane fluidity [82].

The node-of-Ranvier experiences an earlier onset of rupture and mechanoporation under deformation. In contrast, the myelin sheath exhibits a greater resilience to rupture, withstanding higher strain levels before failure. The results also suggest a direct relationship between Young’s modulus and rupture strain (shown in Fig. S5). However, further data and studies are needed to reliably confirm this. Furthermore, the deformation rate and rupture strain appear to be directly related, with slower deformations resulting in lower rupture strains. Examining membrane pores indicates that they often form in regions with higher concentrations of PE lipids, while glycolipids are absent near the pores. The distribution of lipids around the pores is driven by the difference in the strength of interactions at the molecular level. These interactions have been shown to affect local membrane tension that would increase the susceptibility of the membrane to pore formation [83].

By coupling MD and FE simulation results, the injury threshold at cellular level has been identified to be around 10 to 13%. At this axon strain, we observe rupture of Node-of-Ranvier regardless of the axon deformation rate. A direct comparison of the results with available experimental data at cellular and sub-cellular levels is difficult due to different time and length scales of experiments and simulations [84] and lack of experiments on axons at molecular level due to methodological challenges, the complexity of the central nervous system (CNS), and the limitations of current experimental models [85, 86, 87, 88]. Nevertheless, an experimental study conducting concussion to swine animals showed widespread disruption of the node-of-Ranvier and loss of Nav1.6 ion channels in the brain white matter, which is in line with our findings about the earlier rupture of the node-of-Ranvier compared to other parts of axolemma [89]. Interestingly, MD studies on a plasma membrane showed a strain rupture of 0.36, which is 22% lower than the strain rupture of the node-of-Ranvier [15], supporting the importance of properly accounting for lipid composition, tail length, and lipid saturation levels in the description of the cellular membrane.

At cellular level, the specific threshold for axonal strain in the brain is influenced by various factors, including the location of the injury, the direction and magnitude of the forces applied, and the individual characteristics of the brain tissue. Nevertheless, experiments using animal models and computational studies have provided some insights into the biomechanical thresholds associated with axonal injury. For example, animal studies have shown that strains ranging from 10% to 20% can lead to axonal injury in the brain [90]. Another study on the embryos’ neurons showed that strains more than 10% would result in a total break of axonal bundles [28]. Another study that applied 10% Lagrangian strain with the rate of 20 /s to organotypic hippocampal slice cultures showed moderate brain damage that was detected 4 days after the injury but not 2 days after the injury [91]. In contrast to our results, in an experiment on the guinea pig optic nerve done by Bain and Meaney, the range of critical strain for axonal damage was shown to be between 0.14 to 0.34 with a strain rate of 30 to 60 /s, below which no nerve was injured [92]. These studies are used for validation of our results based on the cascading nature of DAI that would lead to further injuries in the damaged cell and also propagate over the brain tissue over time. While some impairments can manifest within minutes of impact, other morphological changes become evident on larger scales days after the injury [93]. Experimental observations in the mentioned studies are assumed to stem from the damage to axons after the impact. However, some differences in the observed experimental injury thresholds and the calculated ones might be due to the ability of the cells to repair themselves when the injury is not severe, which would lead to neuronal impairments at the cellular level within hours after the injury [94]. The difference in injury thresholds might also be due to mechanical differences between animal and human axons or because the axon model that is used in this study is isolated, while previous studies have shown that the axons can withstand larger strains when embedded inside the tissue [39].

It is essential to recognize certain limitations within this study. First, the lack of experimental data at the molecular level on the sub-cellular component of the axon does not allow to build a pure experimental-based model for MD and FE (i.e. lack of axolemma lipidome, or mechanical characterization of the myelinated axon). That results in our membrane model reproducing the lipid composition trend among membrane types, which might diverge from “exact” lipid composition, and using the FE model representing an unmyelinated axon. The material properties assigned to the axon are based on experimental data from the axonal cortex, which may differ from the mechanical properties of the myelinated axon membrane, which has different material properties in different regions.

Another point to consider is that we have used a CG model for the membrane, where a particle represents a group of atoms. The reduction of the degree of freedom affects the entropy of the system (e.g., lipid chain entropy) and might influence membrane rigidity and the timescale of simulations. In addition, Previous studies have shown that the timescale of simulations using coarse-grained modelling and Martini forcefield is 3 to 8 times faster than all-atom (AA) simulations based on the data on diffusion constants [45].

Also, the strain rates observed in experimental studies on brain tissue damage and real-world scenarios are between 10-100 *s*^−1^ [95]. Considering the time mismatch between the CG and AA simulations, the strain rates used for deformation are still at least an order of magnitude faster than experimental rates. However, if an experimental strain rate were used, it is estimated that the simulation would take years to induce lipid rupture, even on a supercomputer with a performance of 5055 TFLOPS on its nodes compared to 0.7 TFLOPs achieved by a node of a typical PC [96]. This would render the process impractical, given the available computational resources and the objectives of the project. Nevertheless, we assumed the simulation rate is already slow enough to present real-world properties of the membrane. In addition, it is not possible to compare the results directly with experiments, as there is no experiment specifically performed on axonal membranes and their behaviour under deformation at length scales investigated in the current study. However, our findings were still in line with previous studies on deriving mechanical properties of other membranes and bilayers. Moreover, coupling the results with FE studies shows that the rupture strain is in line with reported thresholds of axonal injuries in experimental studies, which can validate our results.

Since we use experimental-based molecular models for different regions of the axolemma, we expect that the behaviour under biological conditions and under deformation will reflect the trend observed in reality. The rupture strain can be used as a baseline for the prediction of axonal injury by coupling MD with the FE simulations and the identified threshold can also be used for the prediction of axonal injuries in different parts of the brain at tissue and head levels after impact, in line with previous study conducted by Montanino et al. [97], where an FE analysis of real-world concussion scenario was performed at the brain level, and the strains applied to the tissue in different parts of the corpus callusom (CC) were calculated. By applying the derived strains to tissue-level FE models of the axon embedded inside the matrix, they calculated the maximum local strains of the cortex for each model.

## 5 Conclusion

This study aimed to elucidate the mechanism of axonal injury, specifically focusing on the possible role of the membrane poration under strain. We develop a molecular model of the axolemma, accounting for the available experimental data on lipid-head composition and tail information. Models with different protein concentrations were considered to mimic different axolemma regions. The developed models along with myelin developed by Saeedimasine et al. [16] were deformed until rupture and the mechanical properties were compared. Our findings revealed that myelin exhibited the highest stiffness among axonal membranes and showed a larger rupture strain and stress at the point of pore formation. In contrast, the node-of-Ranvier had the lowest rupture strain and stiffness and counted as the most vulnerable part of the axonal membrane. However, the tension and stress at the pore were not significantly different at different protein concentration, suggesting that these properties are primarily influenced by the lipid composition of the membrane rather than the protein concentration.

We combined the lipid-bilayer MD simulations with FE simulations of the axon to get an better understanding of the mechanism behind axonal injury and its initial trigger and shed light on the behaviour of sub-cellular elements of the axons during an impact. We get the rupture strain for each membrane model from MD and the axonal strain at which local membrane mechanoporation occurs from FE simulations and then compare the injury threshold with experimental findings. The simulation results indicate that the mechanoporation of the membrane occurs in the range of 10 to 13 % of axonal strain, which aligns with the experimental data and supports the hypothesis that mechanoporation might trigger axonal injury. In addition, the study highlights how combining FE and MD approaches can explore complex multi-scale phenomena that cannot be investigated by a single method.

## Author contributions

**Maryam Majdolhosseini:** Methodology, Validation, Formal analysis, Data Curation, Investigation, Resources, Visualization, Writing – original draft. **Svein Kleiven:** Supervision, Resources, Writing – review and editing, Funding acquisition. **Alessandra Villa:** Conceptualization, Methodology, Supervision, Resources, Writing – review and editing, Funding acquisition.

## Declaration of Competing Interest

The authors declare no conflict of interest.

## Acknowledgements

The computational resources were provided by the National Academic Infrastructure for Supercomputing in Sweden (NAISS) through the Center for High-Performance Computing (PDC) at KTH under Projects NAISS 2024/22-601, SNIC 2021/5-459, SNIC 2022/5-645, and SNIC 2021/5-406. We would like to acknowledge Jing Gong and Johan Hellsvik at the KTH PDC Center for their assistance with the installation of the necessary software on the PDC resources.

## Funding

This study was supported by research funds from Swedish Research Council Grants VR-2020-04496.

## Notes

### Competing Interest Statement

The authors have declared no competing interest.

### Summary of Updates

The whole text was reorganized for more clarity. The statements describing the methods in the results section were deleted in case of repetition or moved to the methods section in case they were mentioned for the first time. For example, the detailed lipid composition of the axolemma and multi-scale approach are now reported in the methods section. In addition, the results and discussion parts are separated to improve the clarity of the manuscript.

## References

[1] G. T. Manley, A. I. R. Maas, Traumatic brain injury: an international knowledge-based approach, Jama 310 (5) (2013) 473–474.

[2] M. E. Komlosh, D. Benjamini, E. B. Hutchinson, S. King, M. Haber, A. V. Avram, L. A. Holtzclaw, A. Desai, C. Pierpaoli, P. J. Basser, Using double pulsed-field gradient MRI to study tissue microstructure in traumatic brain injury (TBI), Microporous and Mesoporous Materials 269 (2018) 156–159.

[3] D. H. Smith, D. F. Meaney, W. H. Shull, Diffuse axonal injury in head trauma, The Journal of head trauma rehabilitation 18 (4) (2003) 307–316.

[4] S. S. Humble, L. D. Wilson, L. Wang, D. A. Long, M. A. Smith, J. C. Siktberg, M. F. Mirhoseini, A. Bhatia, S. Pruthi, M. A. Day, others, Prognosis of diffuse axonal injury with traumatic brain injury, The journal of trauma and acute care surgery 85 (1) (2018) 155.

[5] B. Hille, Ionic channels in excitable membranes. Current problems and biophysical approaches, Biophysical journal 22 (2) (1978) 283–294.

[6] J. S. Trimmer, K. J. Rhodes, Localization of voltage-gated ion channels in mammalian brain, Annu. Rev. Physiol. 66 (2004) 477–519.

[7] J. Ritchie, R. B. Rogart, Density of sodium channels in mammalian myelinated nerve fibers and nature of the axonal membrane under the myelin sheath., Proceedings of the National Academy of Sciences 74 (1) (1977) 211–215.

[8] I. Ogiwara, H. Miyamoto, N. Morita, N. Atapour, E. Mazaki, I. Inoue, T. Takeuchi, S. Itohara, Y. Yanagawa, K. Obata, others, Nav1. 1 localizes to axons of parvalbumin-positive inhibitory interneurons: a circuit basis for epileptic seizures in mice carrying an Scn1a gene mutation, Journal of Neuroscience 27 (22) (2007) 5903–5914.

[9] A. Duflocq, B. Le Bras, E. Bullier, F. Couraud, M. Davenne, Nav1. 1 is predominantly expressed in nodes of Ranvier and axon initial segments, Molecular and Cellular Neuroscience 39 (2) (2008) 180–192.

[10] T. Harayama, H. Riezman, Understanding the diversity of membrane lipid composition, Nature Reviews Molecular Cell Biology 19 (2018) 281. URL 10.1038/nrm.2017.138 http://10.0.4.14/nrm.2017.138

[11] W. Jiang, Y. C. Lin, Y. L. Luo, Mechanical properties of anionic asymmetric bilayers from atomistic simulations, Journal of Chemical Physics 154 (22) (2021). doi:10.1063/5.0048232.

[12] A. Rasouli, Y. Jamali, E. Tajkhorshid, O. Bavi, H. N. Pishkenari, Mechanical properties of ester- and ether-DPhPC bilayers: A molecular dynamics study, Journal of the Mechanical Behavior of Biomedical Materials 117 (2021). doi:10.1016/j.jmbbm.2021.104386.

[13] D. Drabik, G. Chodaczek, S. Kraszewski, M. Langner, Mechanical Properties Determination of DMPC, DPPC, DSPC, and HSPC Solid-Ordered Bilayers, Langmuir 36 (14) (2020). doi:10.1021/acs.langmuir.0c00475.

[14] M. Saeedimasine, A. Montanino, S. Kleiven, A. Villa, Role of lipid composition on the structural and mechanical features of axonal membranes: a molecular simulation study, Scientific reports 9 (1) (2019) 1–12.

[15] A. Montanino, M. Saeedimasine, A. Villa, S. Kleiven, Localized axolemma deformations suggest mechanoporation as axonal injury trigger, Frontiers in neurology 11 (2020) 25.

[16] M. Saeedimasine, A. Montanino, S. Kleiven, A. Villa, Elucidating axonal injuries through molecular modelling of myelin sheaths and nodes of Ranvier, Frontiers in Molecular Biosciences 8 (2021) 669897.

[17] M. C. Oliveira, M. Yusupov, A. Bogaerts, R. M. Cordeiro, Molecular dynamics simulations of mechanical stress on oxidized membranes, Biophysical Chemistry 254 (2019) 106266.

[18] G. H. DeVries, C. J. Zmachinski, The Lipid Composition of Rat CNS Axolemma-enriched Fractions, Journal of Neurochemistry 34 (2) (1980). doi:10.1111/j.1471-4159.1980.tb06613.x.

[19] G. H. DeVries, W. J. Zetusky, C. Zmachinski, V. P. Calabrese, Lipid composition of axolemma-enriched fractions from human brains., Journal of lipid research 22 (2) (1981) 208–216.

[20] G. H. DeVries, W. Payne, R. G. Saul, Composition of axolemma-enriched fractions isolated from bovine CNS myelinated axons, Neurochemical Research 6 (5) (1981). doi:10.1007/BF00964391.

[21] M. N. Rasband, W. B. Macklin, J. A. Benjamins, Myelin Structure and Biochemistry, Basic Neurochemistry: Principles of Molecular, Cellular, and Medical Neurobiology: Eighth Edition (2012) 180–199doi:10.1016/B978-0-12-374947-5.00010-9.

[22] N. U. Olsson, A. J. Harding, C. Harper, N. Salem, High-performance liquid chromatography method with light-scattering detection for measurements of lipid class composition: Analysis of brains from alcoholics, Journal of Chromatography B: Biomedical Applications 681 (2) (1996). doi:10.1016/0378-4347(95)00576-5.

[23] H.I. Ingólfsson, M. N. Melo, F. J. van Eerden, C. Arnarez, C. A. Lopez, T. A. Wassenaar, X. Periole, A. H. de Vries, D. P. Tieleman, S. J. Marrink, Lipid Organization of the Plasma Membrane, Journal of the American Chemical Society 136 (41) (2014) 14554–14559. doi:10.1021/ja507832e. URL https://doi.org/10.1021/ja507832e

[24] M. H. Magdesian, F. S. Sanchez, M. Lopez, P. Thostrup, N. Durisic, W. Belkaid, D. Liazoghli, P. Grütter, D. R. Colman, Atomic force microscopy reveals important differences in axonal resistance to injury, Biophysical Journal 103 (3) (2012). doi:10.1016/j.bpj.2012.07.003.

[25] A. N. Hellman, B. Vahidi, H. J. Kim, W. Mismar, O. Steward, N. L. Jeon, V. Venugopalan, Examination of axonal injury and regeneration in micropatterned neuronal culture using pulsed laser microbeam dissection, Lab on a Chip 10 (16) (2010). doi:10.1039/b927153h.

[26] A. Selfridge, D. Preece, V. Gomez, L. Z. Shi, M. W. Berns, A model for traumatic brain injury using laser induced shockwaves, in: K. Dholakia, G. C. Spalding (Eds.), Optical Trapping and Optical Micromanipulation XII, Vol. 9548, SPIE, 2015, p. 95480P. doi:10.1117/12.2189724. URL https://doi.org/10.1117/12.2189724

[27] R. S. Chung, J. A. Staal, G. H. McCormack, T. C. Dickson, M. A. Cozens, J. A. Chuckowree, M. C. Quilty, J. C. Vickers, Mild axonal stretch injury in vitro induces a progressive series of neurofilament alterations ultimately leading to delayed axotomy, Journal of Neurotrauma 22 (10) (2005). doi:10.1089/neu.2005.22.1081.

[28] Y. Li, C. Li, C. Gan, K. Zhao, J. Chen, J. Song, T. Lei, A precise, controllable in vitro model for diffuse axonal injury through uniaxial stretch injury, Frontiers in Neuroscience 13 (OCT) (2019). doi:10.3389/fnins.2019.01063.

[29] D. K. Cullen, V. N. Vernekar, M. C. LaPlaca, Trauma-induced plasmalemma disruptions in three-dimensional neural cultures are dependent on strain modality and rate, Journal of neurotrauma 28 (11) (2011) 2219–2233.

[30] M. C. LaPlaca, V. M.-Y. Lee, L. E. Thibault, An in vitro model of traumatic neuronal injury: loading rate-dependent changes in acute cytosolic calcium and lactate dehydrogenase release, Journal of neurotrauma 14 (6) (1997) 355–368.

[31] M. C. LaPlaca, M. C. Lessing, G. R. Prado, R. Zhou, C. C. Tate, D. Geddes-Klein, D. F. Meaney, L. Zhang, Mechanoporation is a potential indicator of tissue strain and subsequent degeneration following experimental traumatic brain injury, Clinical Biomechanics 64 (2019) 2–13.

[32] Pettus Edward H, C. W. Christman, M. L. Giebel, J. T. Povlishock, Traumatically induced altered membrane permeability: its relationship to traumatically induced reactive axonal change, Journal of neurotrauma 11 (5) (1994) 507–522.

[33] R. J. Cloots, J. A. Van Dommelen, T. Nyberg, S. Kleiven, M. G. Geers, Micromechanics of diffuse axonal injury: Influence of axonal orientation and anisotropy, Biomechanics and Modeling in Mechanobiology 10 (3) (2011). doi:10.1007/s10237-010-0243-5.

[34] M. Kazempour, M. Baniassadi, H. Shahsavari, Y. Remond, M. Baghani, Homogenization of heterogeneous brain tissue under quasi-static loading: a visco-hyperelastic model of a 3D RVE, Biomechanics and Modeling in Mechanobiology 18 (4) (2019). doi:10.1007/s10237-019-01124-6.

[35] A. Montanino, S. Kleiven, Utilizing a structural mechanics approach to assess the primary effects of injury loads onto the axon and its components, Frontiers in neurology 9 (2018) 643.

[36] M. D. Tang-Schomer, V. E. Johnson, P. W. Baas, W. Stewart, D. H. Smith, Partial interruption of axonal transport due to microtubule breakage accounts for the formation of periodic varicosities after traumatic axonal injury, Experimental Neurology 233 (1) (2012). doi:10.1016/j.expneurol.2011.10.030.

[37] A. T. N. Vo, M. A. Murphy, P. K. Phan, T. W. Stone, R. K. Prabhu, Molecular dynamics simulation of membrane systems in the context of traumatic brain injury, Current Opinion in Biomedical Engineering (2023) 100453.

[38] M. F. Horstemeyer, R. K. Prabhu, Multiscale Biomechanical Modeling of the Brain, Academic Press, 2021.

[39] A. Montanino, M. Saeedimasine, A. Villa, S. Kleiven, Axons embedded in a tissue may withstand larger deformations than isolated axons before mechanoporation occurs, Journal of biomechanical engineering 141 (12) (2019).

[40] D. Fitzner, J. M. Bader, H. Penkert, C. G. Bergner, M. Su, M.-T. Weil, M. A. Surma, M. Mann, C. Klose, M. Simons, Cell-type-and brain-region-resolved mouse brain lipidome, Cell reports 32 (11) (2020) 108132.

[41] H.I. Ingólfsson, T. S. Carpenter, H. Bhatia, P.-T. Bremer, S. J. Marrink, F. C. Lightstone, Computational lipidomics of the neuronal plasma membrane, Biophysical journal 113 (10) (2017) 2271–2280.

[42] X. Pan, Z. Li, X. Jin, Y. Zhao, G. Huang, X. Huang, Z. Shen, Y. Cao, M. Dong, J. Lei, others, Comparative structural analysis of human Nav1. 1 and Nav1. 5 reveals mutational hotspots for sodium channelopathies, Proceedings of the National Academy of Sciences 118 (11) (2021) e2100066118.

[43] H. Hu, P. Jonas, A supercritical density of Na+ channels ensures fast signaling in GABAergic interneuron axons, Nature neuroscience 17 (5) (2014) 686–693.

[44] K. T. Wann, Neuronal sodium and potassium channels: structure and function, BJA: British Journal of Anaesthesia 71 (1) (1993) 2–14.

[45] S. J. Marrink, H. J. Risselada, S. Yefimov, D. P. Tieleman, A. H. De Vries, The MARTINI force field: coarse grained model for biomolecular simulations, The journal of physical chemistry B 111 (27) (2007) 7812–7824.

[46] T. A. Wassenaar, H. I. Ingólfsson, R. A. Bockmann, D. P. Tieleman, S. J. Marrink, Computational lipidomics with insane: a versatile tool for generating custom membranes for molecular simulations, Journal of chemical theory and computation 11 (5) (2015) 2144–2155.

[47] D. H. de Jong, G. Singh, W. F. D. Bennett, C. Arnarez, T. A. Wassenaar, L. V. Schafer, X. Periole, D. P. Tieleman, S. J. Marrink, Improved parameters for the martini coarse-grained protein force field, Journal of chemical theory and computation 9 (1) (2013) 687–697.

[48] X. Periole, M. Cavalli, S. J. Marrink, M. A. Ceruso, Combining an elastic network with a coarse-grained molecular force field: Structure, dynamics, and intermolecular recognition, Journal of Chemical Theory and Computation 5 (9) (2009). doi:10.1021/ct9002114.

[49] S. Páll, A. Zhmurov, P. Bauer, M. Abraham, M. Lundborg, A. Gray, B. Hess, E. Lindahl, Heterogeneous paralleliza-tion and acceleration of molecular dynamics simulations in GROMACS, Journal of Chemical Physics 153 (13) (2020). doi:10.1063/5.0018516.

[50] Lindahl, Abraham, Hess v. der Spoel, GROMACS 2020.7 Source code (2 2022). doi:10.5281/zenodo.5938877. URL https://doi.org/10.5281/zenodo.5938877

[51] Lindahl, Abraham, Hess v. der Spoel, GROMACS 2021 Source code (1 2021). doi:10.5281/zenodo.4457626. URL https://doi.org/10.5281/zenodo.4457626

[52] M. Bernetti, G. Bussi, Pressure control using stochastic cell rescaling, The Journal of Chemical Physics 153 (11) (2020) 114107.

[53] G. Bussi, T. Zykova-Timan, M. Parrinello, Isothermal-isobaric molecular dynamics using stochastic velocity rescaling, Journal of Chemical Physics 130 (7) (2009). doi:10.1063/1.3073889.

[54] V. Corradi, E. Mendez-Villuendas, H. I. Ingólfsson, R.-X. Gu, I. Siuda, M. N. Melo, A. Moussatova, L. J. DeGagne, B. I. Sejdiu, G. Singh, others, Lipid–protein interactions are unique fingerprints for membrane proteins, ACS central science 4 (6) (2018) 709–717.

[55] S. Pandiyan, P. V. Parandekar, O. Prakash, T. K. Tsotsis, N. N. Nair, S. Basu, Controlling the sub-molecular motions to increase the glass transition temperature of polymers, Chemical Physics Letters 593 (2014). doi: 10.1016/j.cplett.2013.12.076.

[56] E. A. Evans, R. Waugh, L. Melnik, Elastic area compressibility modulus of red cell membrane, Biophysical Journal 16 (6) (1976). doi:10.1016/S0006-3495(76)85713-X.

[57] W. Rawicz, K. C. Olbrich, T. McIntosh, D. Needham, E. A. Evans, Effect of chain length and unsaturation on elasticity of lipid bilayers, Biophysical Journal 79 (1) (2000). doi:10.1016/S0006-3495(00)76295-3.

[58] G. Saher, B. Brügger, C. Lappe-Siefke, W. Möbius, R.-i. Tozawa, M. C. Wehr, F. Wieland, S. Ishibashi, K.-A. Nave, High cholesterol level is essential for myelin membrane growth, Nature neuroscience 8 (4) (2005) 468–475.

[59] I. Bjorkhem, S. Meaney, Brain cholesterol: long secret life behind a barrier, Arteriosclerosis, thrombosis, and vascular biology 24 (5) (2004) 806–815.

[60] O. S. Andersen, R. E. Koeppe, Bilayer thickness and membrane protein function: An energetic perspective (2007). doi:10.1146/annurev.biophys.36.040306.132643.

[61] B. Alberts, A. Johnson, J. Lewis, D. Morgan, M. Raff, K. Roberts, P. Walter, Molecular Biology of the Cell, New York: Garland Science, 2017. doi:10.1201/9781315735368.

[62] R. Ashkar, M. Nagao, P. D. Butler, A. C. Woodka, M. K. Sen, T. Koga, Tuning Membrane Thickness Fluctuations in Model Lipid Bilayers, Biophysical Journal 109 (1) (2015). doi:10.1016/j.bpj.2015.05.033.

[63] X. F. Jiang, K. Yang, X. Q. Yang, Y. F. Liu, Y. C. Cheng, X. Y. Chen, Y. Tu, Elastic modulus affects the growth and differentiation of neural stem cells, Neural Regeneration Research 10 (9) (2015). doi:10.4103/1673-5374.165527.

[64] L. Picas, F. Rico, S. Scheuring, Direct measurement of the mechanical properties of lipid phases in supported bilayers, Biophysical Journal 102 (1) (2012). doi:10.1016/j.bpj.2011.11.4001.

[65] O. Et-Thakafy, N. Delorme, C. Gaillard, C. Mériadec, F. Artzner, C. Lopez, F. Guyomarch, Mechanical Properties of Membranes Composed of Gel-Phase or Fluid-Phase Phospholipids Probed on Liposomes by Atomic Force Spectroscopy, Langmuir 33 (21) (2017). doi:10.1021/acs.langmuir.7b00363.

[66] N. M. Fonseka, F. T. Arce, H. S. Christie, C. A. Aspinwall, S. S. Saavedra, Nanomechanical Properties of Artificial Lipid Bilayers Composed of Fluid and Polymerizable Lipids, Langmuir 38 (1) (2022). doi:10.1021/acs.langmuir.1c02098.

[67] E. A. Evans, R. Waugh, Osmotic correction to elastic area compressibility measurements on red cell membrane, Biophysical Journal 20 (3) (1977). doi:10.1016/S0006-3495(77)85551-3.

[68] B. Daily, E. L. Elson, G. I. Zahalak, Cell poking. Determination of the elastic area compressibility modulus of the erythrocyte membrane, Biophysical Journal 45 (4) (1984). doi:10.1016/S0006-3495(84)84209-5.

[69] J. H. Adams, D. Doyle, D. I. Graham, A. E. Lawrence, D. R. Mclellan, Microscopic Diffuse Axonal Injury in Cases of Head Injury, Medicine, Science and the Law 25 (4) (1985). doi:10.1177/002580248502500407.

[70] A. E. Jolly, M. Balaeţ, A. Azor, D. Friedland, S. Sandrone, N. S. Graham, K. Zimmerman, D. J. Sharp, Detecting axonal injury in individual patients after traumatic brain injury, Brain 144 (1) (2021). doi:10.1093/brain/awaa372.

[71] F. E. Sherriff, L. R. Bridges, S. Sivaloganathan, Early detection of axonal injury after human head trauma using immunocytochemistry for β-amyloid precursor protein, Acta Neuropathologica 87 (1) (1994). doi:10.1007/BF00386254.

[72] R. M. Venable, F. L. Brown, R. W. Pastor, Mechanical properties of lipid bilayers from molecular dynamics simulation (2015). doi:10.1016/j.chemphyslip.2015.07.014.

[73] B. W. Peeters, A. C. Piët, M. Fornerod, Generating Membrane Curvature at the Nuclear Pore: A Lipid Point of View, Cells 11 (3) (2022). doi:10.3390/cells11030469.

[74] Y. Ha, J. H. Kwon, Effects of lipid membrane composition on the distribution of biocidal guanidine oligomer with solid supported lipid membranes, RSC Advances 10 (38) (2020). doi:10.1039/d0ra03108a.

[75] Y. Ma, K. Poole, J. Goyette, K. Gaus, Introducing membrane charge and membrane potential to T cell signaling (2017). doi:10.3389/fimmu.2017.01513.

[76] R. J. Clarke, Electrostatic switch mechanisms of membrane protein trafficking and regulation (2023). doi:10.1007/s12551-023-01166-2.

[77] K. Hac-Wydro, P. Wydro, The influence of fatty acids on model cholesterol/phospholipid membranes, Chemistry and Physics of Lipids 150 (1) (2007). doi:10.1016/j.chemphyslip.2007.06.213.

[78] S. Chakraborty, M. Doktorova, T. R. Molugu, F. A. Heberle, H. L. Scott, B. Dzikovski, M. Nagao, L. R. Stingaciu, R. F. Standaert, F. N. Barrera, J. Katsaras, G. Khelashvili, M. F. Brown, R. Ashkar, How cholesterol stiffens unsaturated lipid membranes, Proceedings of the National Academy of Sciences of the United States of America 117 (36) (2020). doi:10.1073/pnas.2004807117.

[79] H. P. Duwe, E. Sackmann, Bending elasticity and thermal excitations of lipid bilayer vesicles: Modulation by solutes, Physica A: Statistical Mechanics and its Applications 163 (1) (1990). doi:10.1016/0378-4371(90)90349-W.

[80] N. Kučerka, J. D. Perlmutter, J. Pan, S. Tristram-Nagle, J. Katsaras, J. N. Sachs, The effect of cholesterol on short- and long-chain monounsaturated lipid bilayers as determined by molecular dynamics simulations and X-ray scattering, Biophysical Journal 95 (6) (2008). doi:10.1529/biophysj.107.122465.

[81] D. Marsh, Lateral pressure in membranes (1996). doi:10.1016/S0304-4157(96)00009-3.

[82] J. A. Lundbæk, P. Birn, A. J. Hansen, R. Søgaard, C. Nielsen, J. Girshman, M. J. Bruno, S. E. Tape, J. Egebjerg, D. V. Greathouse, G. L. Mattice, R. E. Koeppe, O. S. Andersen, Regulation of Sodium Channel Function by Bilayer Elasticity: The Importance of Hydrophobic Coupling. Effects of Micelle-forming Amphiphiles and Cholesterol, Journal of General Physiology 123 (5) (2004). doi:10.1085/jgp.200308996.

[83] N. Bavi, C. D. Cox, Y. A. Nikolaev, B. Martinac, Molecular insights into the force-from-lipids gating of mechanosensitive channels (2023). doi:10.1016/j.cophys.2023.100706.

[84] S. Hosmane, A. Fournier, R. Wright, L. Rajbhandari, R. Siddique, I. H. Yang, K. T. Ramesh, A. Venkatesan, N. Thakor, Valve-based microfluidic compression platform: Single axon injury and regrowth, Lab on a Chip 11 (22) (2011). doi:10.1039/c1lc20549h.

[85] J. Kim, M. S. Sajid, E. F. Trakhtenberg, The extent of extra-axonal tissue damage determines the levels of CSPG upregulation and the success of experimental axon regeneration in the CNS, Scientific Reports 8 (1) (2018). doi: 10.1038/s41598-018-28209-z.

[86] J. P. Lezana, S. Y. Dagan, A. Robinson, R. S. Goldstein, M. Fainzilber, F. C. Bronfman, M. Bronfman, Axonal PPARγ promotes neuronal regeneration after injury, Developmental Neurobiology 76 (6) (2016). doi:10.1002/dneu.22353.

[87] S. M. Han, H. S. Baig, M. Hammarlund, Mitochondria Localize to Injured Axons to Support Regeneration, Neuron 92 (6) (2016). doi:10.1016/j.neuron.2016.11.025.

[88] D. Kilinc, J. M. Peyrin, V. Soubeyre, S. Magnifico, L. Saias, J. L. Viovy, B. Brugg, Wallerian-like degeneration of central neurons after synchronized and geometrically registered mass axotomy in a three-compartmental microfluidic chip, Neurotoxicity Research 19 (1) (2011). doi:10.1007/s12640-010-9152-8.

[89] H. Song, P. P. McEwan, K. E. Ameen-Ali, A. Tomasevich, C. Kennedy-Dietrich, A. Palma, E. J. Arroyo, J. P. Dolle, V. E. Johnson, W. Stewart, D. H. Smith, Concussion leads to widespread axonal sodium channel loss and disruption of the node of Ranvier, Acta Neuropathologica 144 (5) (2022). doi:10.1007/s00401-022-02498-1.

[90] A. E. Davis, Mechanisms of traumatic brain injury: Biomechanical, structural and cellular considerations, Critical Care Nursing Quarterly 23 (3) (2000). doi:10.1097/00002727-200011000-00002.

[91] B. Morrison, H. L. Cater, C. D. Benham, L. E. Sundstrom, An in vitro model of traumatic brain injury utilising two-dimensional stretch of organotypic hippocampal slice cultures, Journal of Neuroscience Methods 150 (2) (2006). doi:10.1016/j.jneumeth.2005.06.014.

[92] A. C. Bain, D. F. Meaney, Tissue-level thresholds for axonal damage in an experimental model of central nervous system white matter injury, Journal of Biomechanical Engineering 122 (6) (2000). doi:10.1115/1.1324667.

[93] J. Ma, K. Zhang, Z. Wang, G. Chen, Progress of research on diffuse axonal injury after traumatic brain injury (2016). doi:10.1155/2016/9746313.

[94] M. O. Krucoff, S. Rahimpour, M. W. Slutzky, V. R. Edgerton, D. A. Turner, Enhancing nervous system recovery through neurobiologics, neural interface training, and neurorehabilitation (2016). doi:10.3389/fnins.2016.00584.

[95] Z. Zhou, X. Li, Y. Liu, W. N. Hardy, S. Kleiven, Brain strain rate response: Addressing computational ambiguity and experimental data for model validation, Brain Multiphysics 4 (2023). doi:10.1016/j.brain.2023.100073.

[96] PDC, About the Dardel HPC system (11 2024). URL https://www.pdc.kth.se/hpc-services/computing-systems/about-the-dardel-hpc-system-1.1053338

[97] A. Montanino, X. Li, Z. Zhou, M. Zeineh, D. Camarillo, S. Kleiven, Subject-specific multiscale analysis of concussion: from macroscopic loads to molecular-level damage, Brain Multiphysics 2 (2021). doi:10.1016/j.brain.2021.100027.

