## supplementary material for "Molecular dynamics study of stiffness and rupture of axonal membranes"

Table S1: Lipid composition of the axolemma, including the mole percentage of each lipid type in the leaflets.

| Model | Extracellular leaflet | Cytoplasmic leaflet |
|---|---|---|
| **PC** | **20.50** | **14.54** |
| DPPC 16:0-18:0 | 5.05 | 3.44 |
| POPC 16:0-18:1 | 3.26 | 2.33 |
| DOPC 16:1-18:1 | 10.62 | 7.55 |
| PAPC 16:0/20:4 | 0.84 | 0.67 |
| DAPC 20:4-22:5 | 0.74 | 0.55 |
| **PE** | **10.73** | **27.19** |
| POPE 16:0/18:1 | 3.15 | 7.99 |
| DOPE 16:1-18:1 | 3.15 | 7.99 |
| PAPE 16:0/20:4 | 0.84 | 2.33 |
| DGPE 20:1-22:1 | 1.26 | 3.22 |
| DAPE 20:4-22:5 | 2.31 | 5.66 |
| **PS** | **0.00** | **5.77** |
| PAPS 16:0/20:4 | 0.00 | 0.67 |
| DAPS 20:4-22:5 | 0.00 | 0.44 |
| PRPS 16:0/24:6 | 0.00 | 4.66 |
| **SM** | **3.26** | **1.44** |
| DPSM 18:1/16:0 | 1.47 | 0.67 |
| POSM 18:1/18:1 | 1.16 | 0.55 |
| PNSM 18:1/24:1 | 0.63 | 0.22 |
| **PI** | **0.00** | **7.55** |
| POPI 16:0/18:1 | 0.00 | 5.99 |
| PAPI 16:0/20:4 | 0.00 | 1.55 |
| **Cerebroside** | **17.14** | **0.00** |
| DPGS 16:1/16:0 | 7.68 | 0.00 |
| DBGS 20:1/20:0 | 2.84 | 0.00 |
| PNGS 18:1/24:1 | 2.84 | 0.00 |
| DPG1 18:1/18:0 | 1.89 | 0.00 |
| DPG3 18:1/18:0 | 1.89 | 0.00 |

| | | |
|---|---|---|
| **Sulfatide** | **4.10** | **0.00** |
| PXSU 18:1/24:0 | 1.47 | 0.00 |
| PNSU 18:1/24:1 | 2.63 | 0.00 |
| **CHOL** | **44.27** | **49.28** |
| **Total** | **100.00** | **100.00** |

Table S2: Building and simulation of systems

| Model Name | Building Procedure | Dimension ($nm \times nm \times nm$ ) | Protein Concentration (channels/$\mu m^2$) | Equilibration Time ($\mu$s) | Deformation |
|---|---|---|---|---|---|
| Axolemma (small system) | INSANE bilayer builder | $20 \times 20 \times 20$ | 0 | 25 | No |
| Axolemma (large system) | axolemma (small system) | $38 \times 38 \times 24$ | 0 | 20 | Yes |
| One-protein-axolemma | axolemma (large system) | $46 \times 38 \times 19$ | 588 | $30^a$ | Yes |
| Small axolemma with protein | axolemma (small system) | $20 \times 20 \times 20$ | 2519 | 25 | No |
| Node-of-Ranvier | small axolemma with protein | $80 \times 80 \times 20$ | 2519 | $15^a + 11.5^b$ | Yes |

a: Applying position restraint on proteins; b: No position restraint on proteins

Table S3: Material properties assigned to the cortex in FE model and the membrane they represent

| **Cortex representative** | **Material properties** |
|---|---|
| Unmyelinated axon [1] | Lin. Viscoel. $G = 0.0016$ MPa; $\eta = 1 \times 10^5$ Pa $\cdot$ sec |
| Node-of-Ranvier | Lin. Viscoel. $G = 8.9^{-4}$ MPa; $\eta = 5.6 \times 10^4$ Pa $\cdot$ sec |
| Myelin | Lin. Viscoel. $G = 0.0021$ MPa; $\eta = 1.3 \times 10^5$ Pa $\cdot$ sec |

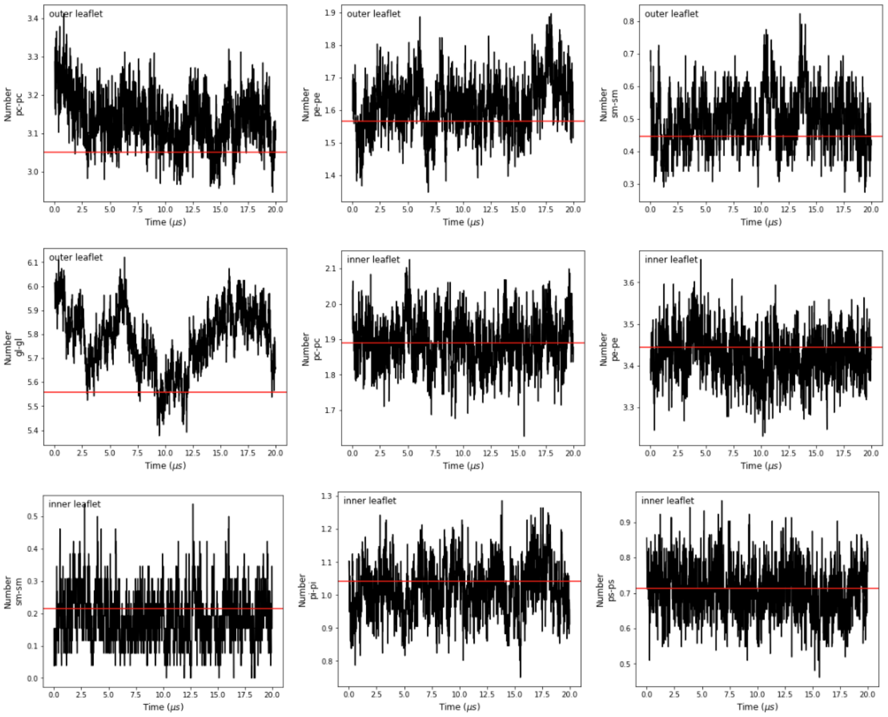


Fig. S1: Number of lipid neighboring within 1.5 nm for the axolemma system of size 40 nm, the average neighboring number in the last 10 $\mu s$ for the system with size of 20 nm is shown in red

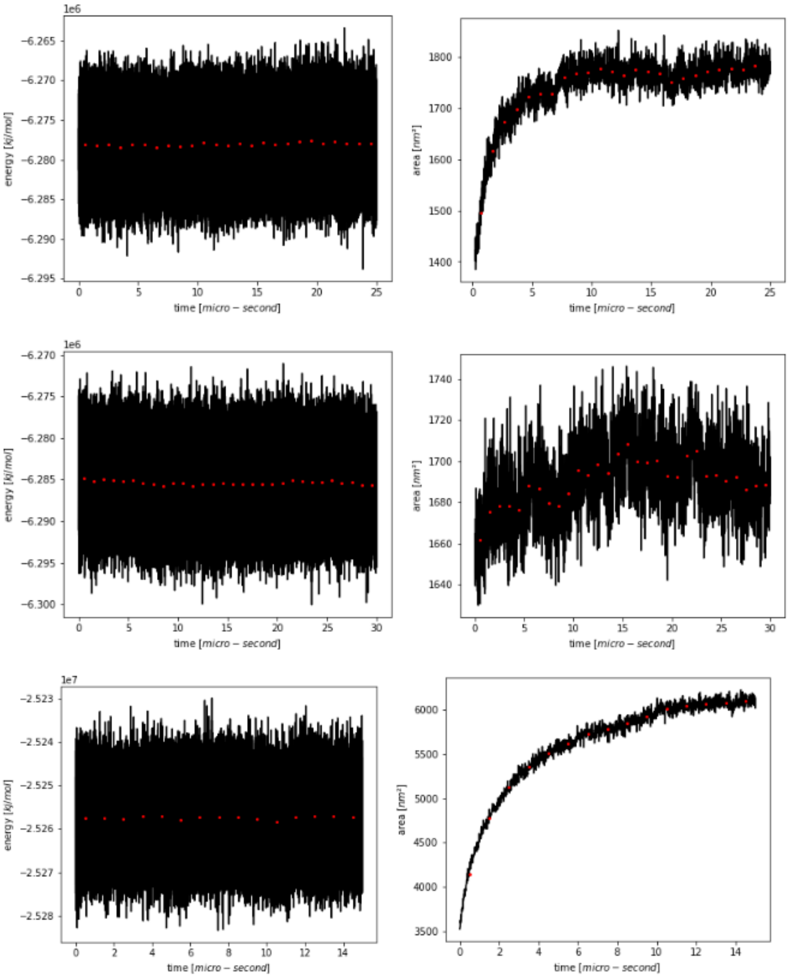


Fig. S2: energy and surface area of systems in equilibrium, average values are taken every 1 $\mu s$ for systems a)axolamme, b) one-protein-axolemma, c)Node-of-Ranvier

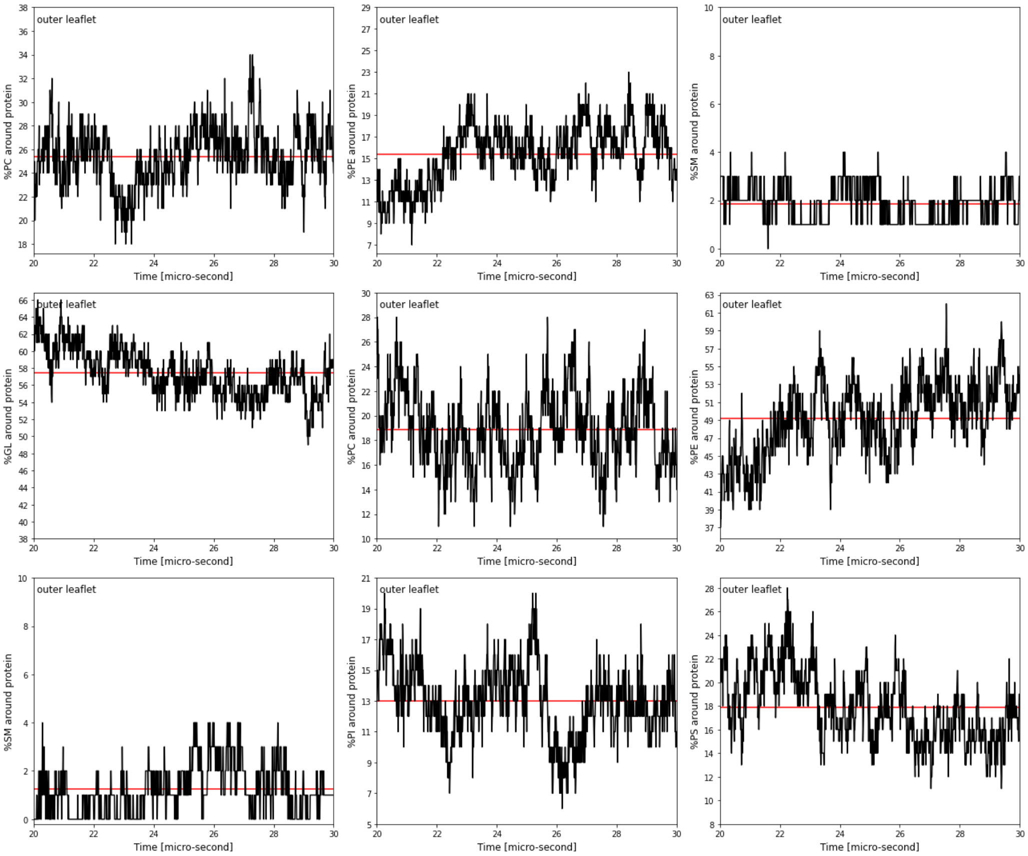


Fig. S3: Protein neighboring for the one-protein-axolemma with position restraint on the protein (black error-bars) at the last 10 $\mu s$ of equilibration and comparison with average protein neighboring for the same system without position restraint on protein at the last 10 $\mu s$ of simulation.

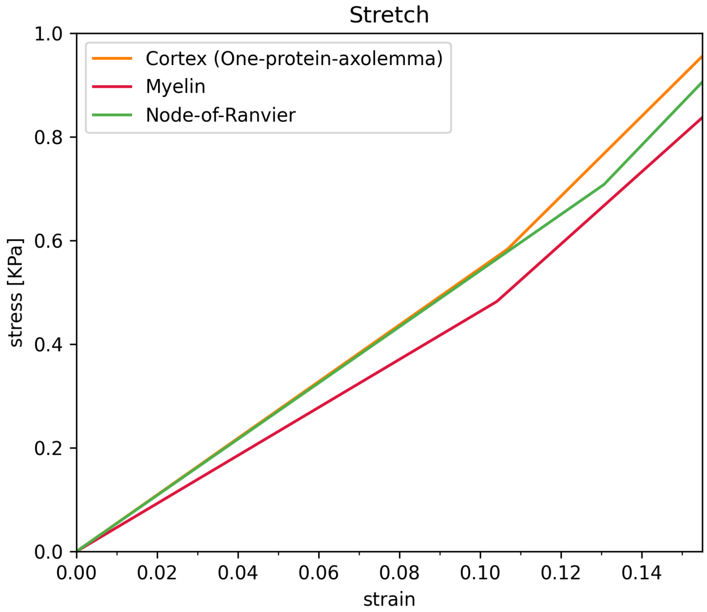


Fig. S4: Maximum local strain vs. axonal strain in uniaxial deformation of the axon when changing the material properties of the cortex.

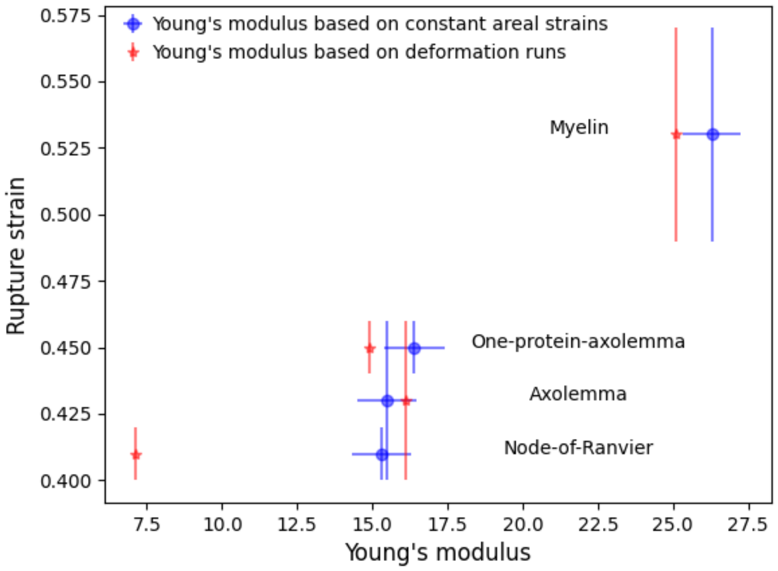


Fig. S5: relationship between Young's modulus and rupture strain. Red dots show rupture strain vs. Young's modulus derived from stress-strain curves, and the blue dots plot rupture strain vs. Young's modulus derived using simulations at constant pressure and areal strain condition, $NP_zA$. Rupture strains are provided based on the simulations with a deformation rate of $10^{-7}$ $nm/ps$

### 1. References


[1] A. Montanino, M. Saeedimasine, A. Villa, S. Kleiven, Localized axolemma deformations suggest mechanoporation as axonal injury trigger, Frontiers in neurology 11 (2020) 25
